# SARS-CoV-2 Viral Pseudoparticles Preferentially Infect Ectoderm In Human Embryonic Tissues

**DOI:** 10.1101/2025.06.27.662024

**Authors:** Ann Song, Prue Talbot

## Abstract

Early stages in human development are difficult to study in pregnant women. We used a “disease-in-a-dish” model to investigate SARS-CoV-2 infection of human embryonic stem cells and the three germ layers. Ectoderm had a significantly higher infection than the other cell types. This was due to: (1) the use of two entry pathways by the ectoderm (fusion and endocytosis), (2) high levels of TMPRSS2 in the ectoderm, and (3) a much-reduced ectodermal glycocalyx, which facilitated viral attachment to the ACE2 receptor. Our findings provide strong evidence that cells in young postimplantation human embryos are susceptible to SARS-CoV-2 infection, which could be embryo lethal or teratogenic in surviving embryos. The high level of infection in ectoderm is a concern as its derivatives may also be affected by SARS-CoV-2. Future clinical work should investigate the functioning of the nervous system in infants born to mothers who had COVID-19 during pregnancy.

**Highlights:**

SARS-CoV-2 pseudoparticles infected human embryonic stem cells, endoderm, mesoderm and ectoderm
Ectoderm was significantly more susceptible to infection than the other three cell types
Factors accounting for increased susceptibility and tissue tropism were identified
SARS-CoV-2 virus can adversely affect early stages of human development

## 1 Introduction

During the COVID-19 pandemic, numerous studies were conducted on human-to-human viral transmission (Guo et al., 2020; Rahman et al., 2020), vaccine development (Jackson et al., 2020; Killeen et al., 2023), and drug development (Arbel et al., 2022; Akinosoglou et al., 2022). However, few reports have examined vertical transmission of the SARS-CoV-2 virus from mother to embryo/fetus in pregnant women with COVID-19 (Kotlyar et al., 2021). The first documented case of vertical transmission occurred in a pregnant woman who contracted COVID-19 in her third trimester (Vivanti et al., 2020). Moreover, maternal SARS-CoV-2 infection has been associated with adverse reproductive outcomes (Young et al., 2022), including an increased risk of pre-eclampsia (Mendoza et al., 2020; Villar et al., 2021; Gurol-Urganci et al., 2021; Wei et al., 2021), premature rupture of the membranes (Allotey et al., 2020), Cesarean delivery (Metz et al., 2021; Gurol-Urganci et al., 2021), and miscarriage (Gurol-Urganci et al., 2021).

*In utero* transmission of SARS-CoV-2 virus is well documented during the late third trimester (Vivanti et al., 2020; Hosier et al., 2020), with studies using postpartum fetal specimens demonstrating transplacental transmission of SARS-CoV-2, whereby the virus crosses the placenta to reach the fetus. SARS-CoV-2 virus has been identified using qPCR (amniotic fluid, nasopharyngeal/rectal swab, blood) and immunohistochemistry in the placentas of infected women and in the cord blood of neonates (Vivanti et al., 2020; Hosier et al., 2020). The strongest evidence of vertical transmission comes from a study of a stillborn fetus during the first trimester (Valdespino-Vázquez et al., 2021). In this study, SARS-CoV-2 virus was detected in placental and fetal tissues by qPCR, immunofluorescence microscopy, and electron microscopy. Fetal organs, such as the lungs and kidneys, were permissive to SARS-CoV-2, leading to significant inflammation and organ damage. Another study examined the vertical transmission during early pregnancy in asymptomatic women (Fenizia et al., 2022). They examined 17 SARS-CoV-2 positive pregnant women who voluntarily terminated their pregnancies during the first trimester and found that 30% of their embryos/fetuses and 20% of syncytiotrophoblasts tested positive for SARS-CoV-2 virus, using qPCR and QuantiGene assays. This supports the idea that SARS-CoV-2 can spread to the embryo/fetus through the placental membrane during early pregnancy, raising concerns regarding the potential long-term consequences of fetal development. In contrast, some authors have suggested that vertical transmission is rare (Peng et al., 2020; Bwire et al., 2021); however, these studies may have not accurately captured the optimal timing for positive PCR testing (Li et al., 2024a) or may have focused on incorrect endpoints such as gross anatomical deficits.

The infection potential of human embryos and fetuses by SARS-CoV-2 raises concerns regarding embryonic lethality and teratogenic effects. Other viruses, such as the Zika virus, which causes microcephaly (Brasil et al., 2016), and cytomegalovirus (CMV), which leads to deafness (Carlson et al., 2010), are well-known teratogens. Accumulating evidence suggests that SARS-CoV-2 may also function as a teratogen in humans. Studies examining fetuses of mothers with COVID-19 have reported restricted fetal growth (Gulersen et al., 2020; Bui et al., 2024), decreased fetal movement (Favre et al., 2020), fetal distress (Villar et al., 2021; Wei et al., 2021; Bui et al., 2024; Zia et al., 2024), stillbirths (Gurol-Urganci et al., 2021; Allotey et al., 2020; Li et al., 2024a), preterm births (Chinn et al., 2020; Metz et al., 2021; Gurol-Urganci et al., 2021; Villar et al., 2021; Wei et al., 2021; Li et al., 2024b; Li et al., 2024a; Zia et al., 2024), neonatal mortality (Villar et al., 2021; Bui et al., 2024), admission to neonatal intensive care units (Metz et al., 2021; Villar et al., 2021; Li et al., 2024a), and low birth weight (Metz et al., 2021; Villar et al., 2021; Wei et al., 2021; Li et al., 2024a). Furthermore, some anomalies associated with SARS-CoV-2 may not be immediately apparent at birth, similar to diethylstilbestrol, which causes vaginal cancer many years after *in utero* exposure (Veurink et al., 2005).

First- and third-trimester studies provide robust evidence of vertical transmission, but an 8-week limitation creates a knowledge gap regarding early embryonic stages during this critical developmental period. To address these knowledge gaps, our study aimed to determine if earlier stages in human development become infected with the SARS-CoV-2 virus and if different cell types are equally susceptible to infection. We used a “disease-in-a-dish-model” based on SARS-CoV-2 pseudoparticles to study infection of four cell types found in early stages of human prenatal development (Song et al., 2023a, b) since experiments on pregnant women are generally not ethically acceptable (Lin and Talbot, 2011). Human embryonic stem cells (hESCs), which have the properties of epiblast cells in post-implantation human embryos (Nichols and Smith, 2009; 2011), were used in conjunction with cells of the three germ layers. SARS-CoV-2 pseudoparticles have a fluorescent viral backbone that allows viral incorporation to be observed microscopically and quantified using flow cytometry. This system enabled us to determine the SARS-CoV-2 viral entry mechanism in hESCs and differentiated germ layer cells, identify drugs that prevent infection, and characterize the causes of tissue tropism.

## 2 Results

### 2.1 Differentiation of Germ Layers from H9 hESCs

Directed differentiation of H9 hESCs into ectoderm, endoderm, and mesoderm was performed with the STEMdiff Trilineage Kit. Each germ layer was labeled with a lineage specific marker. None of the germ layer markers were expressed in the undifferentiated hESCs (Figure S1). After differentiation, markers were expressed in the correct germ layer (endoderm: SOX17, mesoderm: NCAM, and ectoderm: PAX6) without cross reaction to other germ layers (Figure S1).

### 2.2 The machinery for SARS-CoV-2 entry (ACE2 and TMPRSS2) is present on hESCs and germ layer cells

ACE2 receptors and the TMPRSS2 protease are part of the infection machinery that enables viral entry by fusion with the host cell plasma membrane (Figure S2) (Zeng et al., 2022; Colaco et al., 2021; Weatherbee et al., 2020). In this model, the SARS-CoV-2 spike protein binds to the ACE2 receptor on the host cell’s plasma membrane, leading to cleavage of the spike protein at the S2 subunit. Cleavage enables the spike protein to fuse with the host cell plasma membrane (Hoffmann et al., 2020; Cai et al. 2020). Viral entry may also occur by endocytosis during which the spike protein binds to the ACE-2 receptor and in the absence of S2 cleavage by TMPRSS2, the virus is incorporated into the cell by endocytosis (Bayati et al., 2021), after which cathepsin L in the endosome cleaves the spike protein enabling its release into the cytosol.

We first determined if and where the viral entry proteins were located on the hESCs and germ layer cells. Host cells were labeled with lectins that bind cell surface glycans (Figure 1). WGA was localized on the surface of hESCs, endoderm, and mesoderm, and Con A was localized on the surface of ectoderm. Images can be enlarged to view cell surface localization in more detail. The four cell types were then labeled with the ACE2 or TMPRSS2 antibodies (Figure 1A-D). The ACE2 and lectin (WGA or Con A) images were merged, and in all four cell types, ACE2 antibody was colocalized with the lectin (Figure 1A-D), indicating it was present in the plasma membrane. However, differences in distribution of ACE-2 were noted among the cell types. In hESCs, ACE2 was mainly distributed as individual proteins (Figure 1A white arrowheads), while some ACE2 was present in small aggregates (Figure 1A orange arrows). In the endoderm and mesoderm, individual ACE2 proteins were present (Figure 1B, C white arrowheads); however, much of the ACE-2 and lectin were localized in numerous large aggregates (Figure 1B, C orange arrows). In the ectoderm, there were individual ACE2 proteins in the plasma membrane (Figure 1D white arrowheads); however, unlike the other germ layers, there was very little aggregation of ACE2 into clusters (Figure 1D orange arrows).

**Figure 1.**
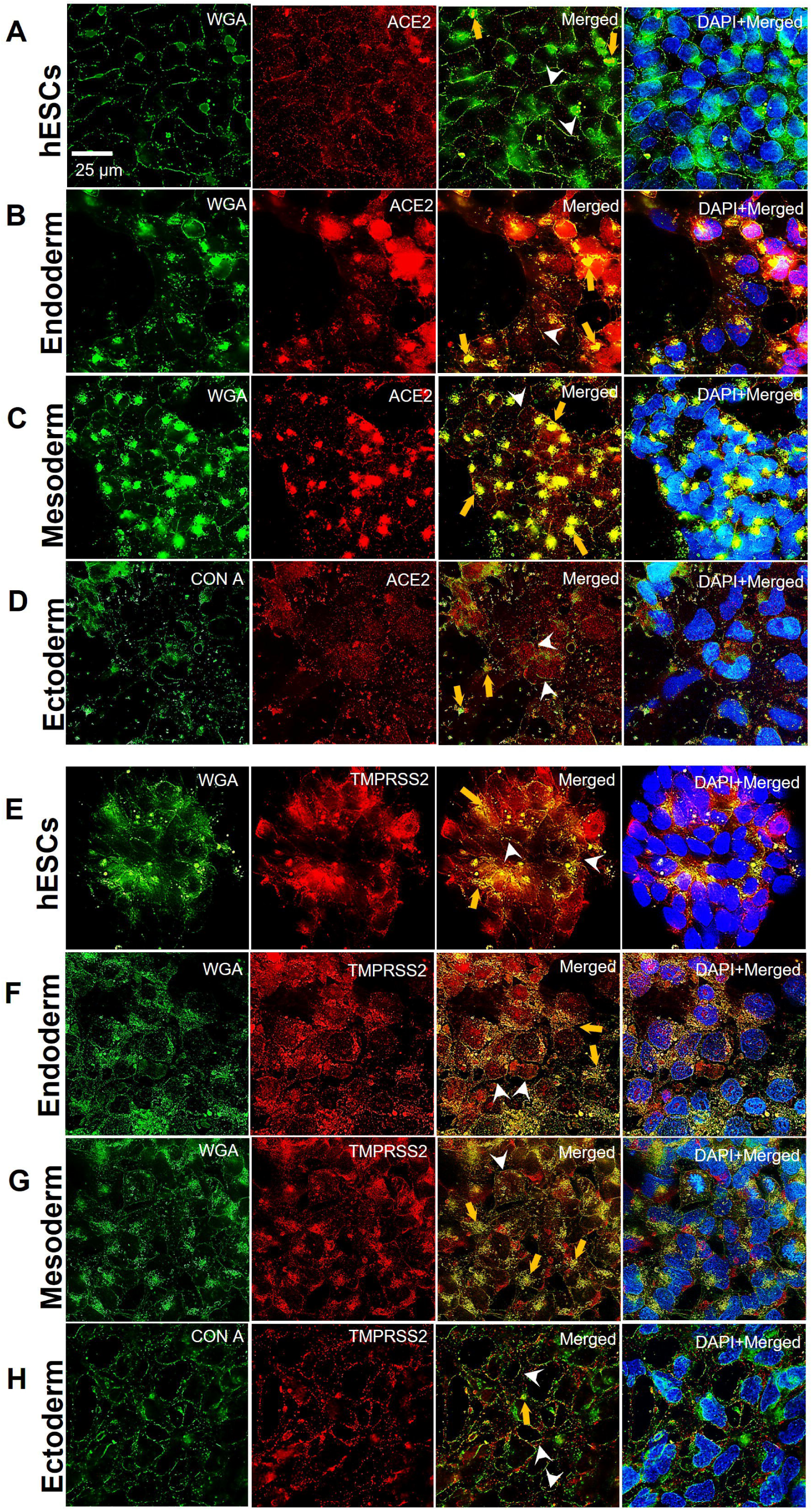
hESCs and their differentiated germ layer cells express the ACE2 receptor and the TMPRSS2 protease. Immunocytochemistry showing ACE2 localization in: (A) hESCs, (B) endoderm, (C) mesoderm, and (D) ectoderm. All samples were labeled with fluorescent lectins (WGA-FITC or ConA-FITC) to verify surface localization of ACE2 and TMPRSS2. Merged images of lectin and ACE2 show colocalization. Immunocytochemistry showing TMPRSS2 localization in: (E) hESCs, (F) endoderm, (G) mesoderm, and (H) ectoderm. Merged images of lectin and TMPRSS2 show colocalization. White arrowheads show individual protein puncta. Orange arrows show protein clusters. Representative images of three independent experiments are shown.

Like ACE2, TMPRSS2 antibody bound to all four cell types and was colocalized with lectin, indicating it was present on the cell surface (Figure 1E-H). In hESCs, individual TMPRSS2 proteins (Figure 1E white arrowheads) and small aggregates were observed (Figure 1E orange arrows). In the endoderm and mesoderm, there were both individual TMPRSS2 proteins in the plasma membrane (Figure 1F, G white arrowheads), and numerous large aggregates of TMPRSS2 and lectin (Figure 1F, G orange arrows). In the ectoderm, there were individual TMPRSS2 proteins (Figure 1H white arrowheads), but aggregates were small and sparse (Figure 1H orange arrows).

These data showed that the four cell types had the machinery needed for SARS-CoV-2 viral entry via the fusion or endocytosis pathways. Both ACE2 and TMPRSS2 were less abundant and less clustered in the endoderm than in the other three cell types.

### 2.3 SARS-CoV-2 pseudoparticles preferentially infect ectoderm

SARS-CoV-2 pseudoparticles were produced using previously published procedures (Song et al., 2023a; Figure 2A). To test the susceptibility of hESCs and the germ layer cells to SARS-CoV-2 infection, a MOI (multiplicity of infection) of 0.1 was used. The mean infection in undifferentiated hESCs was set to 1, and the germ layer groups were compared to this value (Figure 2B). Relative infection varied significantly among the four cell types. Infection levels were similar for the undifferentiated hESCs and endoderm. However, infection was significantly elevated in both mesoderm and ectoderm. The infection in ectoderm was 23-fold greater than in hESCs and 6-fold greater than in mesoderm. These results show that hESCs and germ layer cells are susceptible to SARS-CoV-2 pseudoparticle infection and that ectoderm had a much higher susceptibility than the other three cell types.

**Figure 2.**
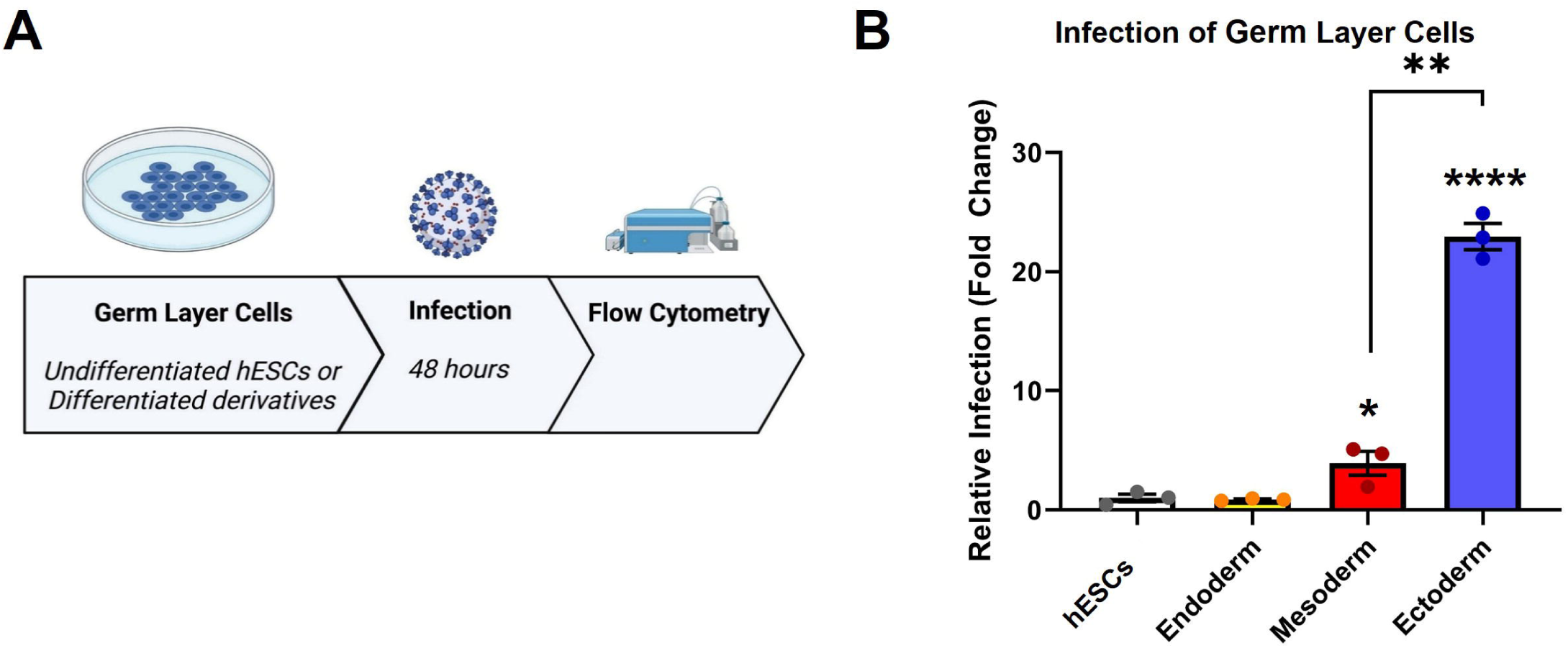
H9 hESCs and their differentiated germ layers were susceptible to SARS-CoV-2 pseudoparticle infection. (A) Flow chart showing the overall infection procedure. (B) Flow cytometry was performed to check the relative infection in the hESCs and germ layer cells. A one-way ANOVA was performed on raw data, and Tukey’s post-hoc test was used to compare differentiated groups to the hESCs. Data are the means ± SEM of three independent experiments. * = p < 0.05, ** = p < 0.01.

### 2.4 SARS-CoV-2 pseudoparticles entered ectoderm via the TMPRSS2 membrane fusion pathway

TMPRSS2 inhibitors (ambroxol, Camostat, aprotinin, Nafamostat) were tested to determine if they could reduce viral infection via the fusion pathway in hESCs and the germ layers. The mean infection in DMSO was set to 100, and the inhibitor groups were compared to this value. Statistical analysis showed that none of the inhibitors were significantly different from the DMSO control in hESCs (Figure 3A), endoderm (Figure 3B), and mesoderm (Figure 3C). However, in ectoderm, aprotinin significantly reduced infection relative to the DMSO control (Figure 3D).

**Figure 3.**
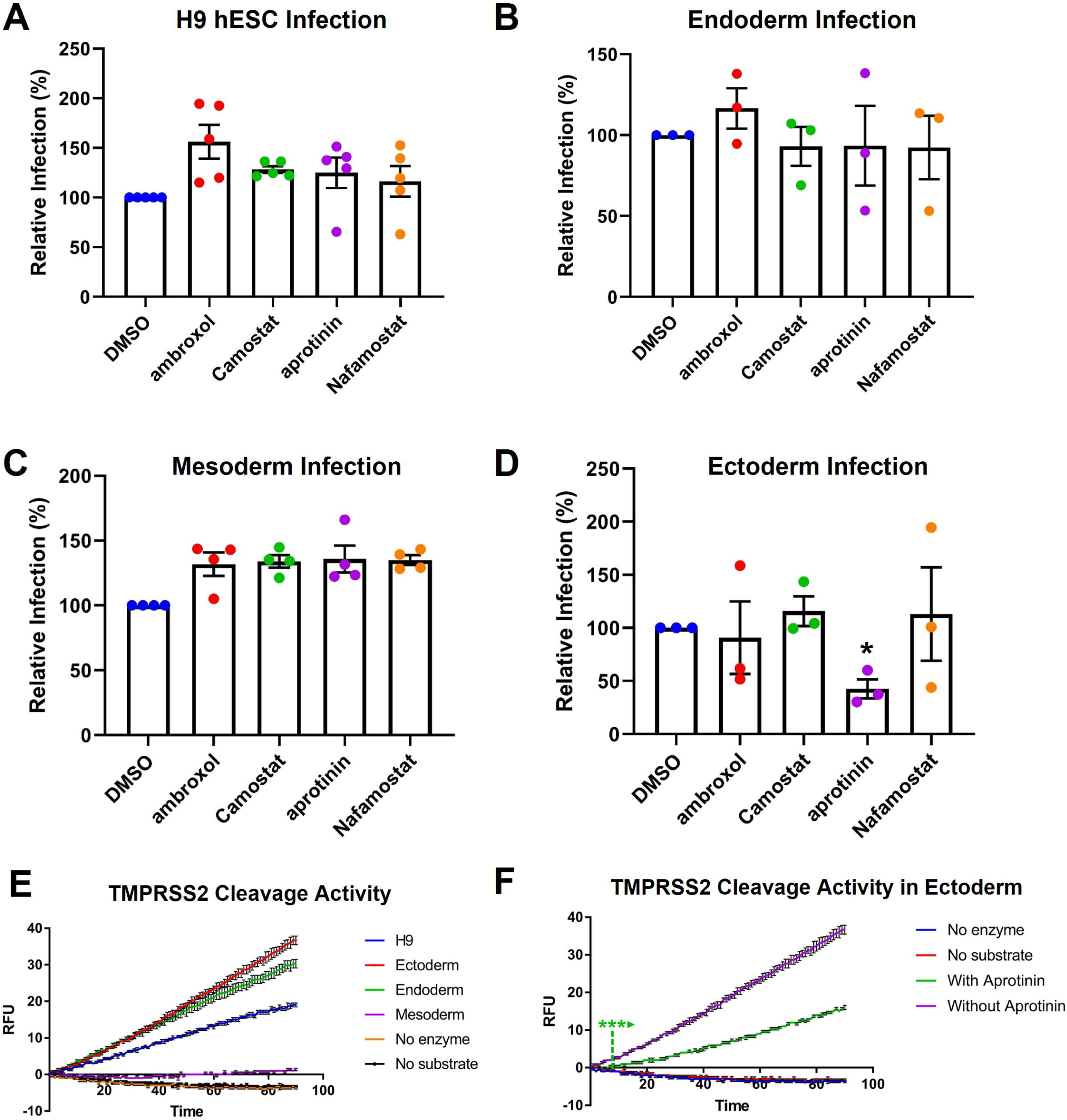
TMPRSS2 inhibitors decreased infection in ectoderm but not in the other cell types. Data are normalized to the DMSO control. SARS-CoV-2 pseudoparticle infection was not significantly affected by TMPRSS2 inhibitors in: (A) hESCs, (B) endoderm, and (C) mesoderm. (D) Aprotinin significantly decreased infection in ectoderm. In (A-D) one-way ANOVAs were performed on raw data, and Dunnett’s post-hoc test was used to compare treated groups to the DMSO control. (E) TMPRSS2 activity of the hESCs and their germ layers. (F) Aprotinin decreased TMPRSS2 cleavage in ectoderm. In (E) and (F), RFU = relative fluorescence units. In (A-F), each group is the mean ± SEM of three independent experiments. * = p < 0.05.

To compare TMPRSS2 activity across cell types, a substrate that fluoresces when cleaved by TMPRSS2 was added to lysates of hESCs and germ layer cells. Lysates from hESCs, ectoderm, and endoderm cleaved the TMPRSS2 substrate when compared to the negative controls (no enzyme, no substrate) (Figure 3E). Ectoderm had the highest enzymatic activity, while mesoderm had almost no activity. Aprotinin significantly reduced protease activity in ectodermal lysates (Figure 3F), supporting its observed reduction of infection (Figure 3D).

### 2.5 SARS-CoV-2 pseudoparticles entered hESCs via endocytosis

Endocytosis inhibitors (Dyngo4a, Pitstop2, nystatin, mβCD, filipin, OcTMAB, MiTMAB) were used to determine if they could reduce infection relative to the DMSO control in hESCs and the germ layer cells. In hESCs, Pitstop2, nystatin, OcTMAB, and MiTMAB reduced infection relative to the DMSO control, but only OcTMAB and MiTMAB were significant (Figure 4A). In the untreated control, dextran-TRITC was taken up by the hESCs and appeared as fluorescent puncta, consistent with uptake by endocytosis (Figure 4B). hESCs treated with OcTMAB or MiTMAB showed little or no uptake of dextran-TRITC.

**Figure 4.**
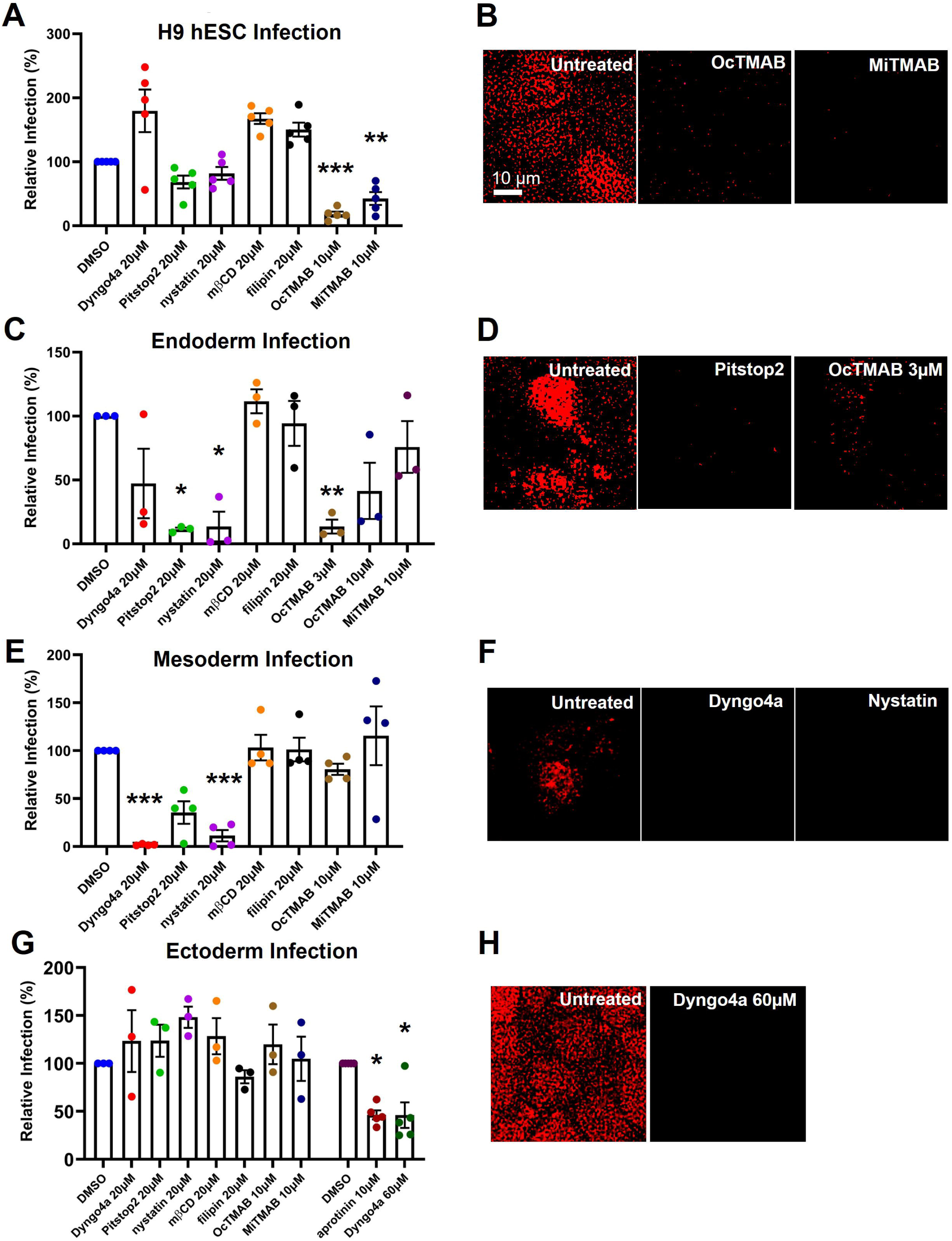
Endocytosis inhibitors decreased infection in all four cell types. (A) OcTMAB and MiTMAB significantly decreased infection in hESCs. (B) OcTMAB and MiTMAB decreased endocytosis of TRITC-conjugated dextran. (C) Pitstop2, nystatin, and OcTMAB (3 µM) significantly decreased infection in endoderm. (D) Pitstop2 and OcTMAB (3 µM) decreased endocytosis of TRITC-conjugated dextran in endoderm. (E) Dyngo4a and nystatin significantly decreased infection in mesoderm. (F) Dyngo4a and nystatin decreased endocytosis of TRITC-conjugated dextran in mesoderm. (G) Dyngo4a (60 µM) significantly decreased infection in ectoderm. (H) Dyngo4a (60 µM) decreased endocytosis of TRITC conjugated dextran. In (A), (C), (E), (G), one-way ANOVAs were performed on the raw infection data, and Dunnett’s post-hoc test was used to compare treated groups to the DMSO control. Data are the means ± SEM of three independent experiments. * = p < 0.05, ** = p < 0.01, *** = p < 0.001.

### 2.6 SARS-CoV-2 pseudoparticles entered the germ layers via endocytosis

In endoderm, Pitstop2 (20 µM), nystatin (20 µM), and OcTMAB (3µM) significantly decreased infection (Figure 4C). Dyngo4a (20 µM) and OcTMAB (10µM) also reduced infection but were not significantly different from the DMSO control. Nystatin treatment caused cell death, so only Pitstop2 and OcTMAB (3µM) were tested for dextran-TRITC uptake (Figure 4D). Endodermal cells treated with Pitstop2 or OcTMAB showed little or no fluorescence compared to the untreated control.

In mesoderm, Dyngo4a (20 µM), Pitstop2 (20 µM), nystatin (20 µM), and OcTMAB (10 µM) decreased infection relative to the DMSO control, but only Dyngo4a and nystatin were significant (Figure 4E). These inhibitors prevented the uptake of dextran-TRITC compared to the untreated control (Figure 4F).

In ectoderm, no effect was observed with endocytosis inhibitors when screened at standard concentrations; however, a single-pass screen (data not shown) at a threefold higher concentration revealed a response from Dyngo4a, prompting three subsequent experiments. Other inhibitors did not respond to increased concentrations. Dyngo4a at a higher concentration (60 µM) significantly reduced infection (Figure 4G). Dextran-TRITC uptake was completely prevented by Dyngo4a (60 µM) compared to the untreated control (Figure 4H).

### 2.7 Endocytosis inhibitors did not alter most germ layer markers

To determine if the drugs that inhibited infection affected germ layer differentiation, qPCR was performed using markers for each cell type. In undifferentiated hESCs, *OCT4* expression was not significantly different in the untreated control and inhibitor treated groups (Figure S3A). In the endoderm, Pitstop2 did not significantly affect *SOX17* expression; however, significant upregulation was observed with OcTMAB (Figure S3B). In the mesoderm, Dyngo4a did not significantly affect *NCAM* expression, but *NCAM* was significantly upregulated with nystatin (Figure S3C). In the ectoderm, the expression of *PAX6* was not significantly different from the control in the aprotinin and Dyngo4a treated groups (Figure S3D).

### 2.7 Cell surface glycosylation affected tropism

The tropism or variation in infectability among the four cell types may have been due in part to the ectoderm having high TMPRRS2 activity and using both the membrane fusion and endocytosis entry pathways, while the other cell types used only endocytosis. To further explore the causes of tropism, we tested the hypothesis that the surface of the ectodermal cells has less glycosylation than the other three cell types, thereby giving the spike protein on viral pseudoparticles better access to the ACE2 receptor on the host cell plasma membrane. To test this hypothesis, the four cell types were first labeled with seven different FITC-conjugated lectins that bind a range of sugar moieties (Figure 5). The amount and pattern of glycosylation were different for each cell type. SBA and DBA did not bind to any of the cell types. In contrast, Con A bound to all cell types. RCA120 bound to all cell types, but the fluorescence was weak in the endoderm and ectoderm. The remaining lectins (WGA, UEA I, PNA) bound to hESCs, endoderm, and mesoderm; but not to the ectoderm. These data show differential glycosylation of the four cell types and support the hypothesis that ectoderm is less glycosylated than the hESCs and the other germ layers.

**Figure 5.**
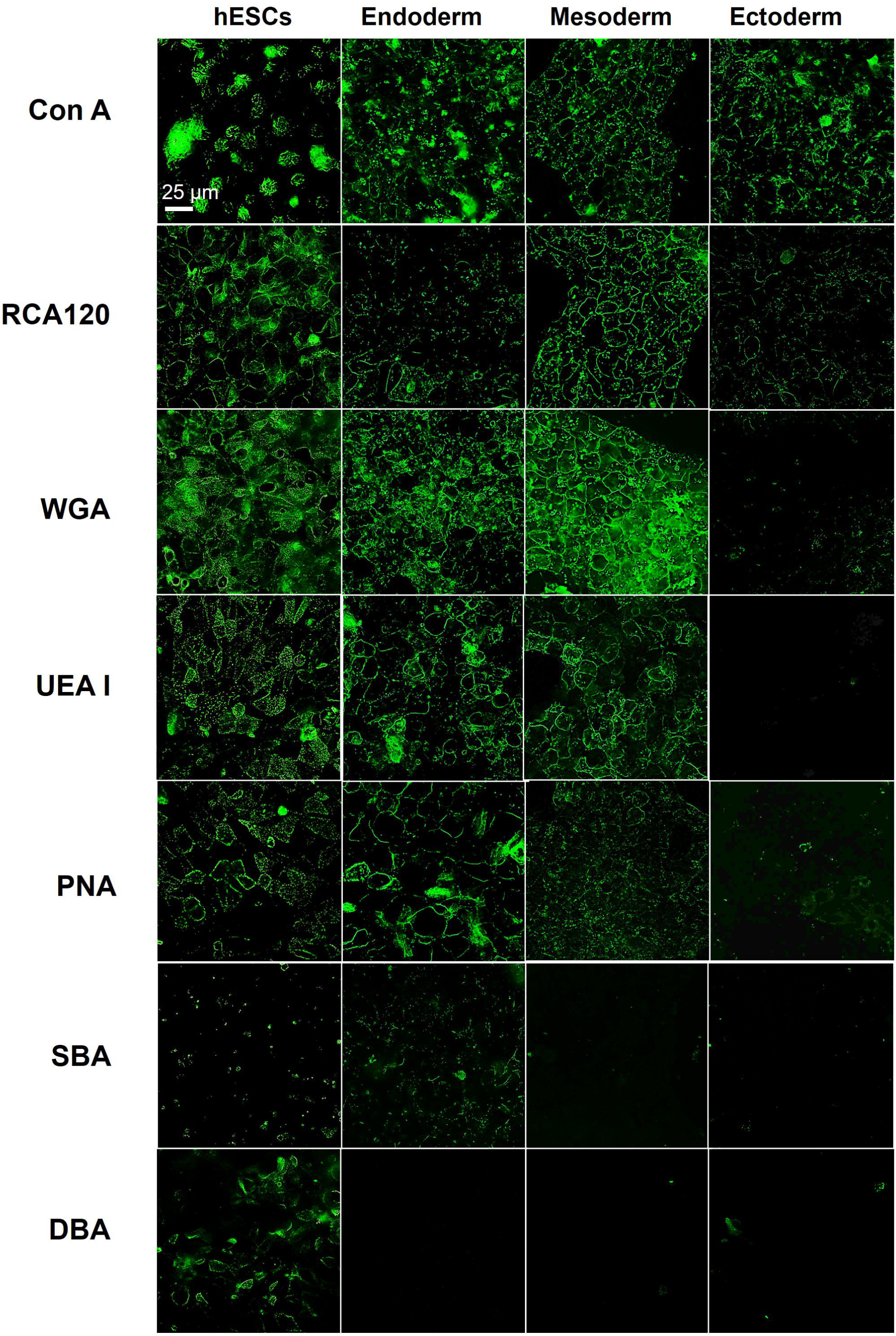
hESCs and the three germ layers showed differential glycosylation when labeled with FITC-conjugated lectins. Various FITC-lectins (ConA, RCA120, WGA, UEA I, PNA, SBA, DBA) were used to label hESCs and the germ layer cells. Of the seven lectins, only ConA bound to the ectoderm, while the other three cell types were well labeled by five (ConA, RCA120, WGA, UEA1 and PNA) of the seven lectins.

As a second step to test the above hypothesis, sialic acid, which binds WGA, was enzymatically removed from cell surfaces with neuraminidase, and SARS-CoV-2 pseudoparticle infection was compared to untreated controls. Sialic acid was chosen for removal since WGA binding was elevated in hESCs, endoderm, and mesoderm, and not observed in ectoderm (Figure 5). Cells were incubated with 0.006 U/mL or 0.018 U/mL of neuraminidase for up to 360 min (0, 45, 90, 180, and 360 min) to remove surface sialic acid. Neuraminidase treatment decreased WGA-FITC binding to hESCs, endoderm, and mesoderm (Figure 6A-C), indicating the removal of some, but not all, sialic acid groups from the cell surface. WGA binding decreased with longer incubation times (e.g., compare hESCs at 90, 180, and 360 minutes) and with higher enzymatic activity (e.g., compare hESCs treated with 0.006 vs 0.018 U/mL of neuraminidase at 90 minutes) (Figure 6A-C).

**Figure 6.**
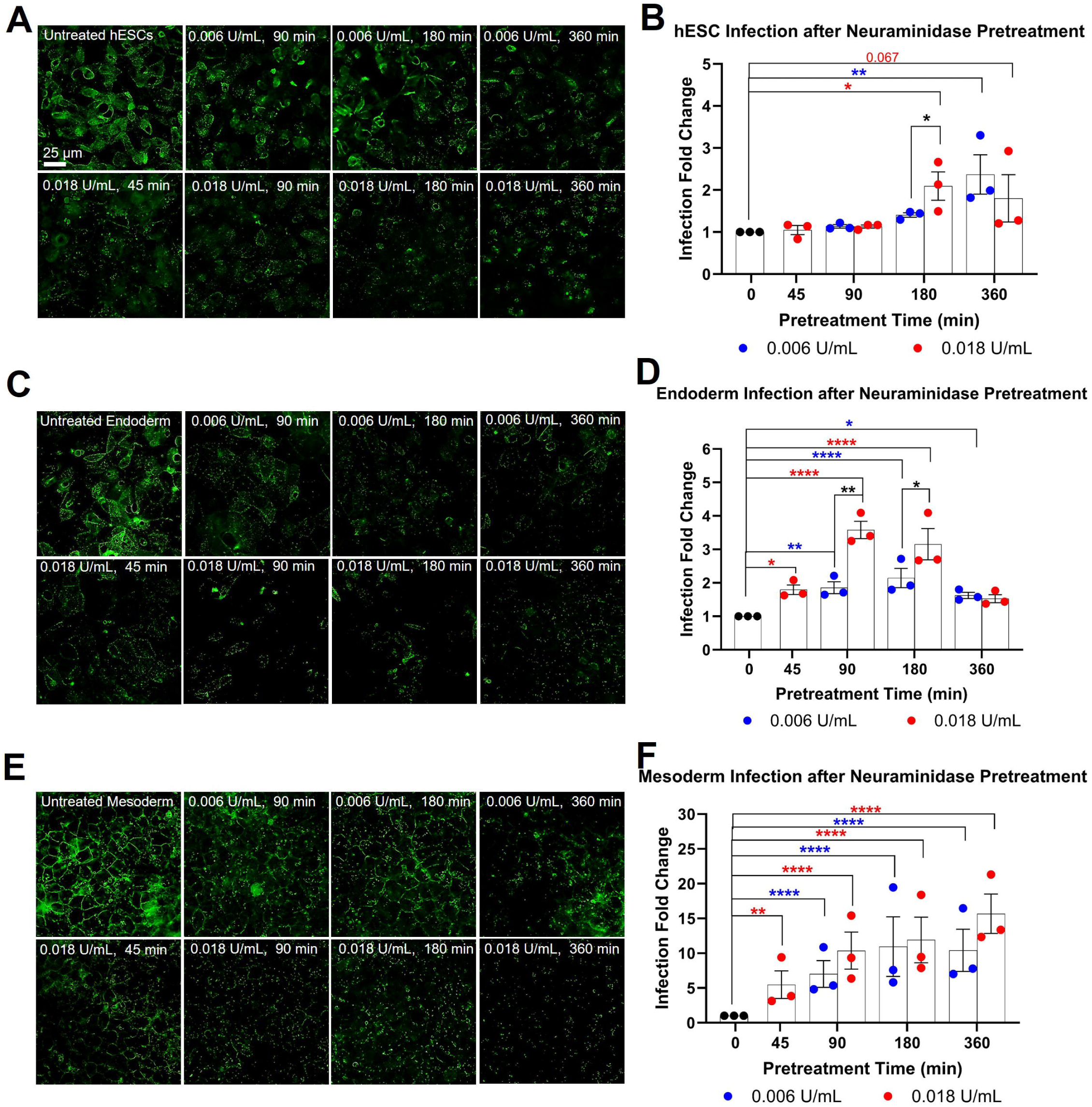
Neuraminidase treatment of hESCs, endoderm, and mesoderm increased infection. (A) H9 hESCs, (C) endoderm, and (D) mesoderm were labeled using FITC-WGA with or without neuraminidase treatment (0.006 U/mL, 0.018 U/mL). Treated cells showed loss of WGA-FITC labeling versus the untreated control. Neuraminidase treatment significantly increased infection in: (B) hESCs, (D) endoderm, and (F) mesoderm, based on one-way ANOVAs on raw infection data followed by Dunnett’s post-hoc tests to compare neuraminidase treated groups to the untreated control. An unpaired two-tailed t-test was used to compare 0.006 U/mL and 0.018 U/mL groups against each other. Data are the means ± SEM of three independent experiments. * = p < 0.05, ** = p < 0.01, **** = p < 0.0001. Blue asterisk = one-way ANOVA significance for 0.006 U/mL of neuraminidase; Red asterisk = one-way ANOVA significance for 0.018 U/mL of neuraminidase; Black asterisk = t-test significance for 0.006 U/mL and 0.018 U/mL comparisons.

To determine if removal of sialic acid would increase infection of hESCs, cells were pretreated with neuraminidase (0.006 U/mL, 0.018 U/mL) for various periods of time before infecting them with SARS-CoV-2 pseudoparticles. In hESCs, a significant increase in infection was observed in the group pretreated for 180 min with 0.018 U/mL of neuraminidase, and a similar increase in infection occurred in the group pretreated for 360 min with 0.006 U/mL of neuraminidase (Figure 6B). In the groups pretreated for 180 min with neuraminidase, 0.018 U/mL produced greater infection than 0.006 U/mL. The maximum increase in infection for the hESCs treated with neuraminidase was slightly over 2-fold, which was not as high as the 23-fold increase observed in the ectoderm (Figure 2B). Together, the decrease in WGA staining correlated with an increase in infection in hESCs.

Endoderm was very susceptible to neuraminidase treatment. WGA labeling was decreased at all times and by both concentrations of neuraminidase (Figure 6C), and infection was increased following pretreatment with both concentrations of neuraminidase (Figure 6D). Pretreatment with 0.006 U/mL of neuraminidase increased infection significantly in all groups. Pretreatment with 0.018 U/mL of neuraminidase significantly increased infection in the 45, 90, 180 min groups; however, in the 360 min group, infection was not significantly different than the control, probably because some cell death occurred. In groups pretreated with neuraminidase for 90 min and 180 min, 0.018 U/mL produced significantly greater infection than 0.006 U/mL. Infection of deglycosylated mesoderm was 2 - 3.5 times greater than in untreated controls.

For mesoderm, neuraminidase treatment decreased WGA-FITC staining in all treated groups (Figure 6E). Pretreatment with 0.006 U/mL of neuraminidase decreased WGA binding as time of pretreatment increased, while pretreatment with 0.018 U/mL of neuraminidase noticeably decreased WGA binding even at shorter pretreatment time. Infection of mesoderm was significantly increased as time of pretreatment with neuraminidase increased, with the higher concentration producing a larger effect faster (Figure 6F). Pretreatment with 0.006 U/mL gave a 10-fold maximum increase in infection in both the 180 min and 360 min groups. Pretreatment with 0.018 U/mL of neuraminidase gave a 16-fold maximum increase in infection, which was close to the ectoderm infection levels (∼23-fold) (Figure 2B). Overall, in both the endoderm and mesoderm, a positive correlation between the loss of WGA staining and an increase in infection was observed, with the mesoderm being especially responsive to removal of sialic acid.

## 3 Discussion

The information on the vertical transmission of SARS-CoV-2 in pregnant women is limited (Chaudhry et al., 2023; Kotlyar et al., 2021; Vivanti et al., 2020; Hosier et al., 2020) and comes mainly from observational and epidemiological studies that deal with the late stages of pregnancy (Overton et al., 2022; Kim and Kim, 2023). Infection early in development is a concern as these stages are often the most sensitive to chemicals and infectious agents, and if the embryo survives, the consequences will likely be more severe than when infection occurs at a later stage (Moore and Persaud, 2024). *In vitro* models are ideally suited for identification of viruses that are embryo lethal or teratogenic (Talbot, 2008; Talbot and Lin, 2011a,b; Lin et al., 2021). In this study, we used a “disease-in-a-dish” model to characterize the susceptibility of hESCs and the three germ layers to infection by SARS-CoV-2 viral pseudoparticles (Song et al., 2023a, b). Our data support the idea that SARS-CoV-2 virus can infect early stages in human embryonic development and could be embryo lethal or teratogenic. While all four cell types were infected by SARS-CoV-2, ectoderm, which gives rise to most of the nervous system and epidermis of the skin, had the highest susceptibility, which was likely due to: (1) its use of both the membrane fusion and endocytosis entry pathways, in contrast to the other three cell types which used only endocytosis; (2) the high level of TMPRSS2 activity in the ectoderm, which would facilitate entry via the membrane fusion pathway; and (3) the low level of glycosylation on ectodermal cells, which would facilitate contact between the viral spike protein and the ACE2 receptor on the host cell’s plasma membrane (Figure 7).

**Figure 7.**
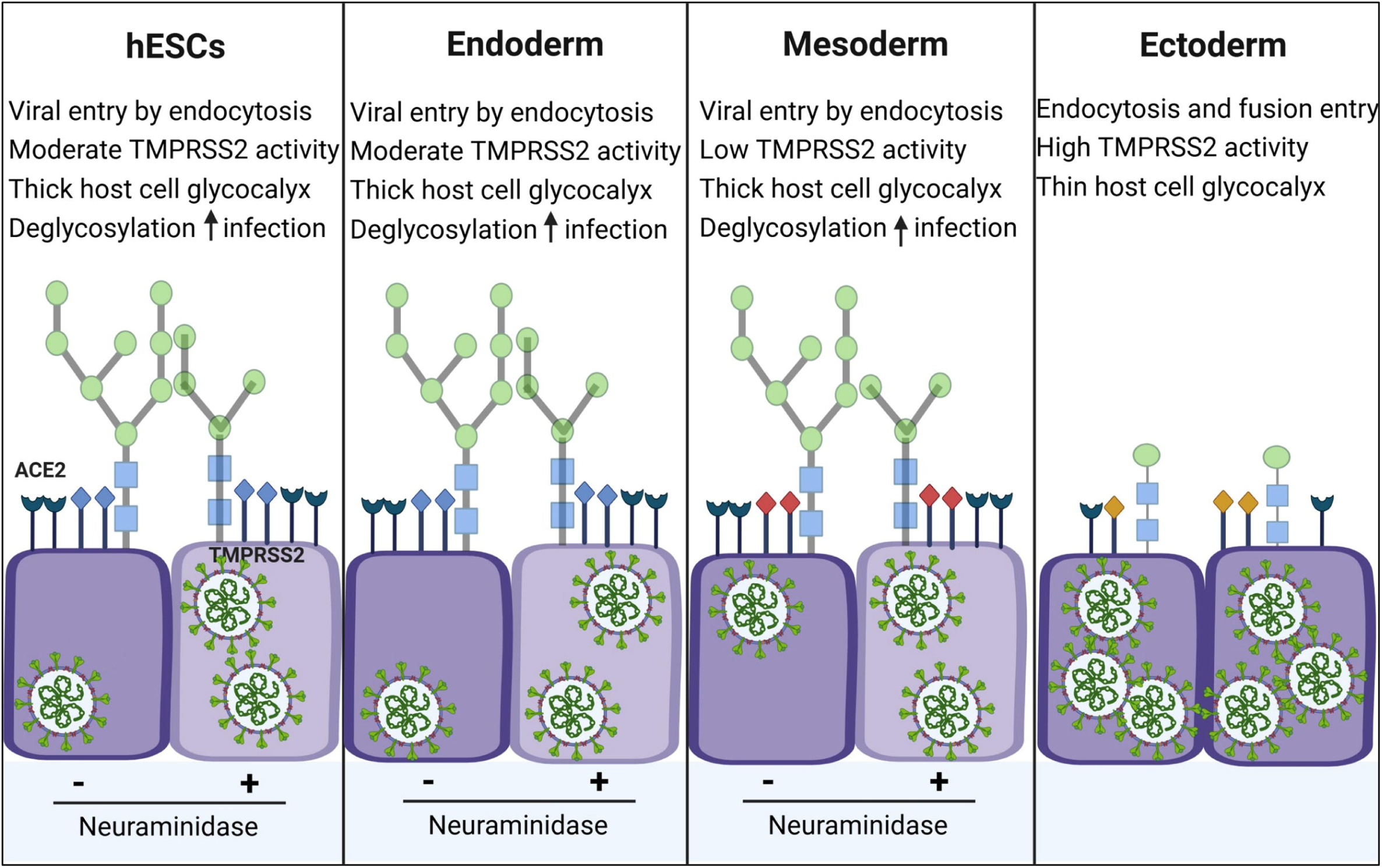
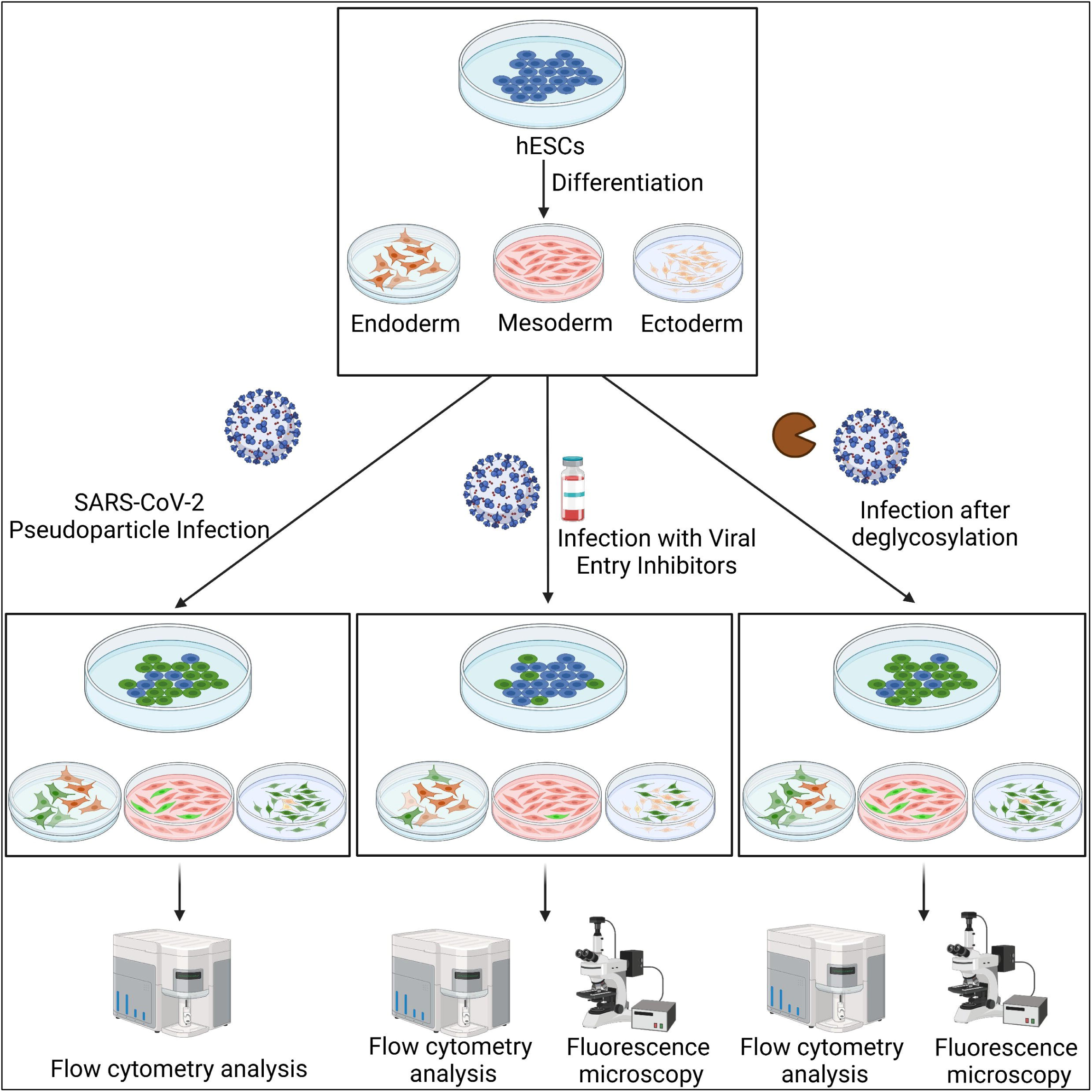
Entry pathways and reasons for tropism in hESCs and germ layer cells. Greater infection of the ectoderm was likely due to three factors: (1) hESCs, endoderm, and mesoderm used only the endocytosis-based pathway for entry into the host cell, while ectoderm used both pathways; (2) TMPRSS2 activity was highest in ectoderm, which likely facilitated its infection by the viral fusion pathway; (3) hESCs, endoderm and mesoderm had heavier glycosylation than ectoderm, and removal of sialic acid with neuraminidase, increased the infectability of hESCs, endoderm, and mesoderm.

Membrane fusion involving spike protein, ACE2, and TMPRSS2 is a major entry pathway for the SARS-CoV-2 virus (Cai et al., 2020; Hoffmann et al., 2020). To investigate the membrane fusion viral entry pathway and potential therapeutic interventions, we tested four TMPRSS2 inhibitors that have been previously studied *in vitro* with SARS-CoV-2 virus or administered to COVID-19 patients (Wang et al., 2023; Carpinteiro et al., 2021; Olaleye et al., 2020; Hoffmann et al., 2020; Hoffmann et al., 2021; Li et al., 2021; Lu et al., 2022; Song et al., 2023b). When tested at 10 µM, only aprotinin reduced SARS-CoV-2 infection, it only worked with ectoderm, and it inhibited both infection and TMPRSS2 activity by about 50%. These results show that not all TMPRSS2 inhibitors are effective at the concentration we tested and that the effects of one inhibitor (aprotinin) cannot be generalized to all cell types. In other studies, aprotinin, at a concentration similar to ours (20 µM), effectively inhibited SARS-CoV-2 infection in Calu-3 cells (Bojková et al., 2020; Bestle et al., 2020) and human primary epithelial cells (Bojková et al., 2020). In a clinical trial, inhaled aprotinin significantly reduced viral load in COVID-19 patients with mild to moderate symptoms (Redondo-Calvo et al., 2022). Together, evidence supports the idea that aprotinin inhibits SARS-CoV-2 infection in ectodermal cells and some human adult cells, and it is the most promising of the protease inhibitors for translation.

Results with other TMPRSS2 inhibitors are mixed. Consistent with our data, 10 µM ambroxol produced little to no inhibition of SARS-CoV-2 infection in HEK293T and A549 cells (Wang et al., 2023), while infection of Vero E6 cells was inhibited by concentrations of ambroxol between 0.1 to 10 µM (Olaleye et al., 2020). At higher concentrations (100-1000 µM), significant inhibition was observed in both the HEK293T and A549 cell lines (Wang et al., 2023), but these concentrations may have acted non-specifically by blocking acid sphingomyelinase (Carpinteiro et al., 2021) or stimulating mucus secretion (Zanasi et al., 2017). In human subjects, a retrospective cohort study reported that ambroxol did not decrease mortality in hospitalized COVID-19 patients (Lu et al., 2022), while bromhexine, its parent drug, significantly reduced the rate of ICU admissions, intubation/mechanical ventilation, and mortality in patients with COVID-19 (Ansarin et al., 2020). Collectively, these studies demonstrate that the *in vitro* response to ambroxol is dependent on cell type and inhibitor concentration, while patients with mild, but not severe, cases of COVID-19 may respond to ambroxol.

*In vitro* studies with camostat and nafamostat have also shown variable results. Camostat and nafamostat (20 µM) were effective in lowering SARS-CoV-2 infection in Calu-3 human lung cells (Hoffmann et al., 2020; Hoffmann et al., 2021), and infection was inhibited by 25 µM concentrations in human airway epithelial cells (Li et al., 2021). However, nonspecific effects, such as inhibiting enteropeptidase activity (Sun et al., 2020), reducing cytokine release (Hoffmann-Wrinkler et al., 2020), and inhibiting blood coagulation (Qian et al., 2023), may occur at these concentrations. In our study, embryonic cells were not responsive to 10 µM camostat or nafamostat. Interestingly, 5 µM was inhibitory in human lung sections (Hoffmann et al., 2021), consistent with our finding that the effective concentration of a protease inhibitor can vary in different tissues (Song et al., 2023b). Two patient cases reported that nafamostat relieved their early developed symptoms (Doi et al., 2020; Takahashi et al., 2021); however, it was not effective in hospitalized patients (Hoffmann-Winkler et al., 2020; Gunst et al., 2021), which is consistent with our conclusion on ambroxol that patients with moderate symptoms may be responsive.

Taken together, the main conclusions from our protease inhibitor data are: (1) only ectoderm used the membrane fusion entry pathway, which may have contributed to its higher infectability; (2) aprotinin was the only TMPRSS2 inhibitor that blocked infection in ectoderm; (3) infection of endoderm, which had high TMPRSS2 activity, was not blocked by any of the protease inhibitors; and (4) if protease inhibitors are used therapeutically in the future, their efficacy may vary with cell type and concentration.

The SARS-CoV-2 virus can also enter host cells by endocytosis, which occurs by both clathrin and caveolae-mediated mechanisms (Bayati et al., 2021). In mink lung epithelium, which has an inactive form of TMPRSS2, endocytosis was the only entry pathway (Song et al., 2023b). In our study, each of the four cell types used the endocytosis entry pathway, and the specific inhibitors that worked varied among the cell types. Pitstop2, Dyngo4a, OcTMAB, MiTMAB inhibited the clathrin pathway, while Nystatin, Dyngo4a, OcTMAB, MiTMAB inhibited the caveolae pathway (Figure 4S).

Pitstop2 (20 uM) inhibited infection in hESCs, endoderm, and mesoderm, with the strongest effects observed in endoderm. Our data agree with studies that demonstrated the effectiveness of Pitstop2 at concentrations similar to ours (10, 12.5, and 15 µM) in A549 cells, human endothelial cells, and HEK293T cells, respectively (Qu et al., 2024; Qian et al., 2021; Bayati et al., 2021). Low (20 µM) and higher (60 µM) concentrations of Pitstop2 had no effect on the ectoderm (data not shown), while efficacy has been reported at a higher concentration (50 µM) in human renal cells (Somova et al., 2024). When testing at high concentration, it is important to consider the potential for nonspecific effects, such as disruption of the nuclear permeability barrier leading to cytotoxicity (Liashkovich et al., 2015). Together, these findings suggest that the cellular response to Pitstop2 is dependent on both the cell type and concentration. Despite its potential as demonstrated in our study and other preclinical studies, clinical trials are essential to establish their safety and efficacy in COVID-19 patients.

Nystatin (20 µM) resulted in a dual effect: it inhibited infection in the mesoderm, yet it was cytotoxic in the endoderm, inducing cell death. Unexpectedly, nystatin enhanced infection of VeroE6 cells (Nguyen et al., 2024). Given its well-documented nonspecific antifungal effects (Baek et al., 2013), hESCs and ectoderm were not tested at high concentrations. The lack of clinical studies further demonstrates that nystatin is unsuitable for research on SARS-CoV-2. Although nystatin has been predicted to be a good TMPRSS2 inhibitor (Virág et al., 2021), our data strongly indicates that it is not a viable candidate for SARS-CoV-2 treatment. Thus, it is important to focus on more promising alternatives to combat SARS-CoV-2.

OcTMAB and MiTMAB are potent inhibitors of both clathrin- and caveolae-mediated endocytosis, effectively blocking the recruitment of dynamin GTPase to the plasma membrane (Figure 4S). In our study, OcTMAB had significant efficacy when tested with hESCs and endoderm, with the latter requiring a lower concentration for optimal results. hESCs were also affected by MiTMAB, indicating the involvement of dynamin-dependent endocytosis (both clathrin- and caveolae-mediated endocytosis) in SARS-CoV-2 entry in this cell type (Figure 4S). A higher concentration of OcTMAB is not advisable, as its analog (Cetyltrimethylammonium bromide) exhibited embryotoxic and teratogenic effects in pregnant mice when 10 mL/kg was orally administered (Isomaa and Ekman, 1975). Despite the absence of patient trials, our results indicate that OcTMAB and MiTMAB exhibit efficacy, suggesting the urgency for future clinical trials to assess the optimal dosage that maximizes efficacy as a promising therapy while minimizing adverse effects. Therefore, evaluation of these drugs in clinical settings is important for the advancement of patient care.

Dyngo4a, a dynamin inhibitor, did not affect hESCs, but produced variable efficacy across the germ layers. It was moderately effective in endoderm and ectoderm, although the latter required a higher concentration. Dyngo4a completely blocked viral entry in mesoderm, which fits well with the observation that mesoderm did not have TMPRSS2 activity and therefore uses only endocytosis for SARS-CoV-2 entry. This result is corroborated by other studies which reported significant efficacy when employing similar concentrations of 12.5-40 µM with human endothelial and Caco-2 cells (Qian et al., 2021; Sun et al., 2024). The concentration we utilized for the ectoderm (60 µM) was similar to effective concentrations used with human renal cells (50 µM) and HEK293T cells (80 µM) (Somova et al., 2024; Bayati et al., 2021). In our study, 60 µM Dyngo4a reduced both infection and dextran-TRITC uptake, indicating endocytosis was effectively blocked and the observed decrease in infection was not due to nonspecific effects.

The efficacy of the endocytosis inhibitors varied with cell types, as we also observed with TMPRSS2 inhibitors. Our data support the conclusion that the SARS-CoV-2 virus enters hESCs and germ layer cells via endocytosis. Endocytosis inhibitors are understudied as SARS-CoV-2 treatments and lack clinical results compared with TMPRSS2 inhibitors, which have been extensively studied. A combination of aprotinin and Dyngo4a may completely inhibit viral entry, as both reduced infection by 50%, thereby shutting down both entry pathways.

Our data showed that glycosylation on the cell surface of hESCs and germ layer cells may be one of the most important factors influencing SARS-CoV-2 infectability. While entry pathways and TMPRSS2 contribute to this process, glycosylation emerges as the most critical determinant. hESCs, endoderm, and mesoderm exhibited remarkable glycosylation, as demonstrated by their binding to ConA, RCA120, WGA, and UEA I, whereas ectoderm bound only to ConA. This distinction highlights the profound impact of glycosylation on tropism. The removal of sialic acid from hESCs, endoderm, and mesoderm using neuraminidase markedly increased their susceptibility to SARS-CoV-2 pseudoparticles. These results support the conclusion that the complexity and thickness of the glycocalyx are crucial in determining host cell infectability, offering a deeper understanding of SARS-CoV-2 tropism. Importantly, neuraminidase enhanced infection most effectively in the mesoderm, surpassing its effects in the endoderm and hESCs. Because the sugar architecture on each cell type is complex and different (Furukawa et al., 2017), the effects of neuraminidase varied among cell types, and further explains tropism seen in the neuraminidase treated cells. Our glycosylation data are further validated by a computational study, which concluded that the glycocalyx on cells creates a steric hindrance that the SARS-CoV-2 virus must overcome to access the ACE2 receptor (Acosta-Gutierrez et al., 2022). Taken together, our data emphasize the importance of glycosylation in SARS-CoV-2 infectability.

The high susceptibility of the ectoderm to SARS-CoV-2 infection suggests that the embryonic/fetal nervous system could be adversely affected in pregnant women with COVID-19. Since ectodermal infection occurs early in development, the embryo may not survive infection, resulting in embryo lethality. Women infected with COVID-19 may lose their pregnancy before realizing that they were pregnant. If the embryo survives but is damaged by infection, SARS-CoV-2 virus may act as a teratogen leading to abnormal development of the ectodermal derivatives. Other viruses are well-known teratogens. For example, Zika virus causes microcephaly or under development of the cerebral cortex and the cerebellum (Brasil et al., 2016; Phan and Holland, 2021). Congenital CMV is the most common cause of birth defects in the U.S. (Thackeray et al., 2014), leading to low birth weight, seizures, rashes, and microcephaly (Adams Waldorf and McAdams, 2013; Thackeray et al., 2014). CMV can also cause defects during childhood, such as progressive hearing loss (Manicklal et al., 2013; Lim and Lyall, 2017). Evidence linking maternal SARS-CoV-2 infection to congenital anomalies is limited. While a recent systematic review reported increased risks (e.g., preterm birth and low birth weight) in newborns of COVID-19 infected women (El-Atawi et al., 2024), some claim that vertical transmission is unlikely to occur (El-Atawi et al., 2024; Chaubey et al., 2021). Calvert et al. (2023) also looked at SARS-CoV-2 infected pregnant women from 6 weeks preconception to 19 weeks of gestation, but they did not find any congenital anomalies in the nervous system (Odds Ratio = 1.02; neural tube defects, hydrocephaly, microcephaly, arhinencephaly, holoprosencephaly, and agenesis of corpus callosum). While these gross anatomical deficits are easily detectable, embryo loss early in pregnancy or subtle anomalies later in development would have been missed. Like CMV, congenital defects from SARS-CoV-2 infection could be latent at the time of birth and appear during childhood. It is critical to carefully monitor children born to pregnant mothers with COVID-19 to ascertain whether latent neurological deficits or other problems appear as the child develops.

In adults, COVID-19 symptoms are not limited to the respiratory system. Both the central and peripheral nervous systems, derivatives of the ectoderm, are often impaired (Guerrero et al., 2021). For example, during SARS-CoV-2 infection, patients have reported a loss of taste and smell (Saniasiaya et al., 2020; Agyeman et al., 2020; Spudich and Nath, 2022), Guillain-Barre syndrome (an autoimmune condition of the peripheral nerves) (Toscano et al., 2020; Rahimi, 2020), and encephalopathy (a broad term for damage or disease that affects the brain) (Poyiadji et al., 2020; Siahaan et al., 2022). In “long COVID”, reports have included fatigue, “brain fog” (e.g., confusion, poor memory, and poor concentration), shortness of breath, and sleep disorder (Perlis et al., 2022; Alkodaymi et al., 2022). Our data show dramatic tropism in early prenatal development with the ectoderm being highly susceptible to SARS-CoV-2 infection. It will be interesting in future studies to determine if similar tropism exists in adults with ectodermal derivatives being highly susceptible.

In conclusion, the major findings of our study were that: (1) human embryonic cells (epiblast and germ layers) were susceptible to SARS-CoV-2 infection, (2) cell types varied in their infection with ectoderm being the most susceptible; and (3) the increased susceptibility of ectoderm was due to its use of two entry pathways (fusion and endocytosis), higher TMPRSS2 activity, and a much reduced glycocalyx that facilitated interaction of the viral spike protein and ACE2 receptor. Small molecule inhibitors, which lowered the infection level in the four cell types, may be translatable and aid in reducing the risk of vertical transmission of the SARS-CoV-2 virus during pregnancy. The high infection level observed in the ectoderm may forecast developmental problems with the nervous system and skin, and these may not be overt at the time of birth. Physicians should examine newborns, toddlers, and children exposed to SARS-CoV-2 *in utero* for possible developmental issues, including those that affect cognition and the nervous system.

### Limitations of the study

While we tested only one MOI, in actual pregnancies, the number of viral particles reaching the embryo will vary and may affect the outcome. The fluorescent-based peptide substrates are not always specific for their intended target (in this case TMPRSS2) (Lauer-Fields et al., 2001; Zou et al., 2023). Some drugs used to identify the entry pathway have been FDA approved, and several are in clinical trials (Ambroxol, Camostat, and Dyngo4a). While these drugs may eventually be used clinically to prevent infection in pregnant women and their embryos/fetuses, there was not a single drug that blocked infection in all cell types. Future studies could test mixtures of drugs at a range of doses and look for synergy, additivity, and antagonism.

## 4 Experimental procedures

### 4.1 Resource availability

Corresponding author: Further information and requests for resources and reagents should be directed to and will be fulfilled by the corresponding author.

Materials availability: This study did not generate new unique reagents.

Data and code availability: The original contributions presented in the study are included in the article, further inquiries can be directed to the corresponding author.

### 4.2 Tissue Culture and Reagents

H9 hESCs (WiCell, Madison, WI) were grown in complete mTeSR plus culture medium (STEMCELL Technologies, Vancouver, Canada). (Lin and Talbot, 2011; Song et al., 2023) Cells were maintained on Matrigel-coated 6-well plates until they reached approximately 80% confluency. For passaging, cells were gently washed once with phosphate-buffered saline (PBS) and removed from the well with 1 mL of ReLeSR (STEMCELL Technologies, Vancouver, Canada). ReLeSR was immediately removed, and colonies were incubated at 37 °C for 4 min, then 1 mL of mTeSR plus was gently added to the well to collect undifferentiated cells. A 1:10 split ratio was used to passage to a new 6-well plate.

HEK 293T and an ACE2-overexpressing cell line (HEK 293T-ACE2) (ATCC, Manassas, VA) were grown in DMEM with high glucose and 10% FBS (Gibco, Carlsbad, CA). Both cell types were grown in T25 flasks until they reached approximately 90% confluency. For passaging, cells were washed once with PBS, lifted from the flask with 1.25 mL of 0.25% trypsin-EDTA for 3 min at 37 °C, and then centrifuged to remove the supernatant. The pellet was resuspended in complete growth medium.

The following small molecule inhibitors were used to target TMPRSS2: ambroxol (10 µM; TCI Chemicals, Portland, OR), aprotinin (10 µM; Tocris, San Diego, CA), Camostat (10 µM; Sigma-Aldrich, Burlington, MA), Nafamostat (10 µM; TCI Chemicals, Portland, OR). Endocytosis inhibitors that were used included: Dyngo4a (20 µM, Selleckchem, Houston, TX), Pitstop2 (20 µM; Abcam, Cambridge, MA), OcTMAB (10 µM; Abcam, Cambridge, MA), MiTMAB (10 µM; Abcam, Cambridge, MA), mβCD (20 µM; Sigma-Aldrich, Burlington, MA), Nystatin (20 µM; Sigma-Aldrich, Burlington, MA), and Filipin (20 µM; Sigma-Aldrich, Burlington, MA).

### 4.3 Trilineage Differentiation

To simulate early embryonic development, H9 hESCs were differentiated into the three germ layers using the STEMdiff^TM^ trilineage differentiation kit (STEMCELL Technologies, Vancouver, Canada) according to the manufacturer’s protocol. Single cells of H9 were obtained after treatment with gentle cell dissociation reagent (STEMCELL Technologies, Vancouver, Canada). Cells were plated on Matrigel-coated 48-well plates in mTeSR plus medium with ROCK inhibitor (10 µM; Tocris, San Diego, CA): 1 × 10^5^ cells per well for ectoderm and endoderm and 2.5 × 10^4^ for mesoderm and the undifferentiated control. Every 24 hours a medium change of each differentiation medium was performed. Samples were infected with SARS-CoV-2 viral pseudoparticles after 5 days (mesoderm and endoderm) or 7 days (ectoderm and undifferentiated H9) of seeding.

### 4.4 Immunofluorescence Microscopy

The cells were seeded in 8-well chamber slides (Ibidi; Gräfelfing, Germany). After differentiation, the cells were fixed at room temperature in 4% paraformaldehyde in PBS for 15 min and washed with PBS. For ACE2 and TMPRSS2 labeling, cells were treated with 50 mM DTT and 6 M guanidine-HCl for 5 minutes prior to quenching with 100 mM iodoacetamide. For other antibodies, cells were permeabilized with 0.1% Triton X-100 for 10 min. The cells were then blocked with 10% donkey serum. Primary antibodies were diluted in blocking buffer. The following primary antibodies were used: mouse anti-PAX6 (1:60; Developmental Studies Hybridoma Bank, Iowa City, IA), goat anti-SOX17 (1:200; R&D Systems, Minneapolis, MN), mouse anti-NCAM (1:30; Developmental Studies Hybridoma Bank, Iowa City, IA), goat anti-ACE2 (1:200; R&D Systems, Minneapolis, MN), and mouse anti-TMPRSS2 (1:200; Santa Cruz Biotechnology, Dallas, TX).

Following overnight incubation with primary antibody at 4 °C, the cells were washed three times with 0.2% Tween in PBS. Next, the cells were incubated in the dark with Alexa Fluor-conjugated fluorescent secondary antibodies (1:500; Invitrogen, Carlsbad, CA). After washing in PBS, the cells were mounted with diluted Vectashield and DAPI for nuclear staining. A Nikon Eclipse Ti inverted microscope was used to image at 40x. The NIS Elements Software and ImageJ software were used to image and analyze the data.

To establish the location of surface proteins (ACE2, TMPRSS2), fixed cells were labeled with the antibody and fluorescence lectin conjugates (WGA-FITC or Con A-FITC; 1:100, Vector Laboratories, Newton, CA, USA). Antibody staining, mounting, and imaging steps were performed as above. To determine surface glycosylation patterns in the four cell types (hESCs, endoderm, mesoderm, and ectoderm), cells were labeled with various lectins (Con A, DBA, PNA, RCA120, SBA, UEA I, WGA; Vector Laboratories, Newton, CA, USA).

### 4.5 TMPRSS2 Activity Assay

TMPRSS2 fluorogenic substrate, Boc-Gln-Ala-Arg-AMC HCl (2.5 mM, Bachem), was prepared in 50 mM Tris (pH 8) and 150 mM NaCl. After differentiation, H9 hESCs and the differentiated germ layer cells were washed twice with PBS and lysed for 1 min on ice in RIPA buffer. The cells were then sheared with a 21-gauge needle, followed by centrifugation at 3,000 rpm for 5 min at 4 °C. The lysate protein was quantified using the Pierce BCA assay kit (Thermo Scientific, Waltham, MA). 10 µg of protein was added to each reaction well. The fluorogenic substrate was added to each well at a final concentration of 10 µM. Fluorescence intensity was measured at 340/440 nm using a BioTek Synergy HTX, multi-mode microplate reader (Winooski, VT). To validate the inhibitory effects of aprotinin in ectoderm, ectodermal cells were preincubated with aprotinin for 24 h before lysate collection.

### 4.6 Lentiviral Production to Generate SARS-CoV-2 Pseudoparticles

SARS-CoV-2 pseudoparticles were generated as described in our previous work (Song et al., 2023a,b). HEK293T cells were plated with antibiotic-free medium at a density of 7 × 10^6^ cells in a T75 flask and transfected using lentiviral plasmids (Song et al., 2023 a,b) and a Lipofectamine3000 Kit (Invitrogen, Carlsbad, CA) according to the manufacturer’s protocol. After overnight incubation, fresh medium was added to cells supplemented with 1% BSA. Conditioned medium was collected and centrifuged 48 h post-transfection. The supernatant was filtered using a 0.45 µm Acrodisc syringe filter (Cytivia Life Sciences, Marlborough, MA), and the filtrate was mixed with 5x polyethylene glycol (Abcam, Cambridge, MA) and precipitated overnight at 4°C. The lentivirus was collected by centrifugation and the pellet was resuspended in Viral Re-suspension Solution (Abcam, Cambridge, MA). Virus aliquots were stored at -80 °C. Prior to use in experiments, the transduction efficiency of each batch of viruses was tested in HEK 293T-ACE2.

### 4.7 Pseudotyping of human SARS-CoV-2 viral pseudoparticles

H9 hESCs and the differentiated germ layers were infected with SARS-CoV-2 viral pseudoparticles using a multiplicity of infection (MOI) of 0.1. The medium was replaced after overnight incubation of the infected cells. Cells were dissociated with gentle cell dissociation reagent 48 h post-infection and washed three times with 0.5% BSA in PBS. After the final wash, the cells were resuspended in the same wash buffer and analyzed using flow cytometry. A Novocyte flow cytometer (Agilent Technologies, Santa Clara, CA) was used to detect ZsGreen in the FITC channel. The resulting flow cytometer files were analyzed using NovoExpress software. Mock infection was used as a background control. Prior to infection, the cells were preincubated for 2 h with TMPRSS2 inhibitors or endocytosis inhibitors, which were kept in the medium until harvesting for flow cytometry.

### 4.8 Endocytosis Assay

Trilineage differentiation was performed in 8-well chamber slides. The cells were incubated overnight with TRITC-conjugated dextran (Invitrogen, Carlsbad, CA) with and without endocytosis inhibitors. After fixation in 4% paraformaldehyde, mounting was performed using diluted Vectashield with DAPI and images were collected using a Nikon Eclipse inverted microscope.

### 4.9 RNA Extraction and Quantitative Real-Time PCR analysis

RNA was extracted from hESCs and the germ layer cells using the RNeasy Mini Kit (Qiagen, Germantown, MD) according to the manufacturer’s protocol. RT-PCR was performed after generating first-strand cDNAs from 1 µg of total RNA using iScript Reverse Transcription Supermix (Bio-Rad, Hercules, CA), as recommended by the manufacturer. qPCR was performed using the SsoAdvanced Universal SYBR Green Supermix in a BIO-RAD CFX connect cycler (Bio-Rad Laboratories, Hercules, CA). Primer sequences are listed in Table S1. Relative expression analysis is presented as 2^-ΔΔCt^ values normalized to the expression of ACTIN and relative to untreated (negative control) samples. All reactions were performed in triplicates.

### 4.10 Deglycosylation Assay

hESCs, endoderm, and mesoderm were seeded in 48-well plates. After 24 hours, the cells were treated with 0.006 U/mL or 0.018 U/mL of neuraminidase (Sigma-Aldrich, Burlington, MA) in DMEM at 37 °C for variable periods of time (0 min, 45 min, 90 min, 180 min, 360 min). 1 Unit of neuraminidase is defined to liberate 1.0 µM of N-acetyl neuraminic acid per minute at pH 5.0 at 37 °C using bovine submaxillary mucin. After deglycosylation, neuraminidase was quenched with culture medium and cells were washed with PBS.

For microscopy, cells were fixed in 4% paraformaldehyde for 15 minutes at room temperature, followed by WGA-FITC labeling and mounting with Vectashield containing DAPI. For infection, cells were transferred to their respective culture media, and SARS-CoV-2 pseudoparticles were added after the deglycosidase treatments to allow infection. After 48 hours, cells were collected and analyzed using flow cytometry.

### 4.11 Data Analysis and Statistics

For infection and TMPRSS2 activity data, the mean and standard error of the mean for three independent experiments were plotted using GraphPad Prism 10 software (GraphPad, San Diego, CA). For infection data, the mean of the DMSO group was set to 100, and the inhibitor groups were compared to this value. Statistical significance was determined using Minitab Statistics Software (Minitab, State College, PA). When the data were not normally distributed, they were subjected to a Box-Cox or logarithmic transformation, and the data were retested to confirm that they satisfied the analysis of variance (ANOVA) model (normal distribution and homogeneity of variances). Infection analyses were performed using a one-way ANOVA, while TMPRSS2 cleavage analysis was performed using a two-way ANOVA, in which the factors were time and cell type. When the means were significant (p < 0.05), groups treated with inhibitors were compared to each other using Tukey’s post hoc analysis or compared to the control group using Dunnett’s post hoc analysis. To analyze qPCR data, the Kruskal-Wallis nonparametric test was performed followed by Dunn’s post hoc test since the untreated control had no variance. For deglycosylation data at two concentrations of neuraminidase, unpaired two-tailed t-tests were performed using GraphPad Prism 10 software (GraphPad, San Diego, CA) to compare the infection levels against each other.

## 4 Acknowledgments

The graphical abstract, Figure 2A, and Figure 7 were created with a licensed version of BioRender.com. We thank Claudia Osuna and Jack Ona for making some of the batches of viral pseudoparticles. We thank Rattapol Phandthong for optimizing the MOI of some of the batches of viral pseudoparticles. We thank the UCR Stem Cell Core for providing access to the Novocyte Flow cytometer. Supported in part by TRDRP Award #R00RG2620, a Graduate Student Researcher award, Graduate Research Mentorship Program Award, Academic Merit Fellowship, Yvonne Danielson Endowed Graduate Award, and the Dissertation Completion Fellowship Award from the UCR Graduate Division, a Committee on Research grant from the UCR Academic Senate, and the AES Research and Graduate Student Funding Program. The content is solely the authors’ responsibility and does not necessarily represent the official views of the funding agencies.

## 5 Author contributions

A.S. and P.T. contributed to conception, data interpretation, manuscript writing, and editing of the submitted version. P.T. was responsible for project administration and funding acquisition. A.S. contributed the sample preparation, data collection, and processing.

