## Supplementary Material for "SARS-CoV-2 Viral Pseudoparticles Preferentially Infect Ectoderm In Human Embryonic Tissues"

Supplementary Information

**Figure S1**

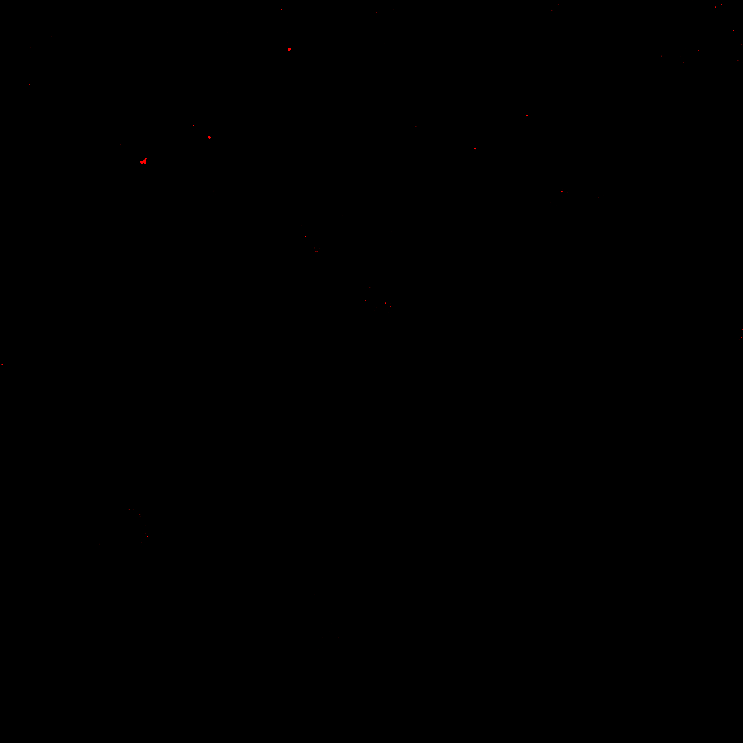

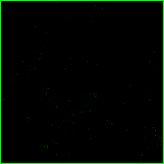

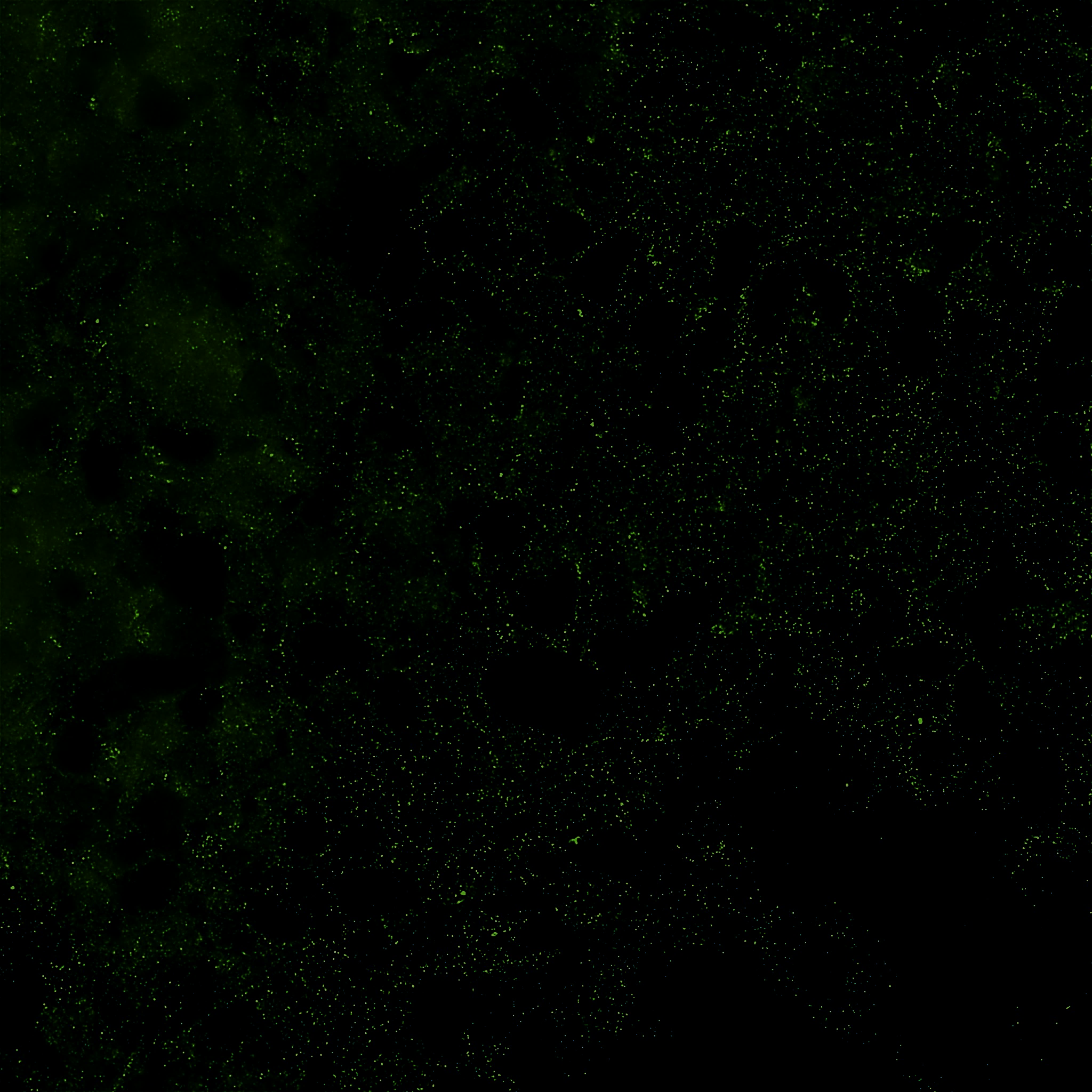

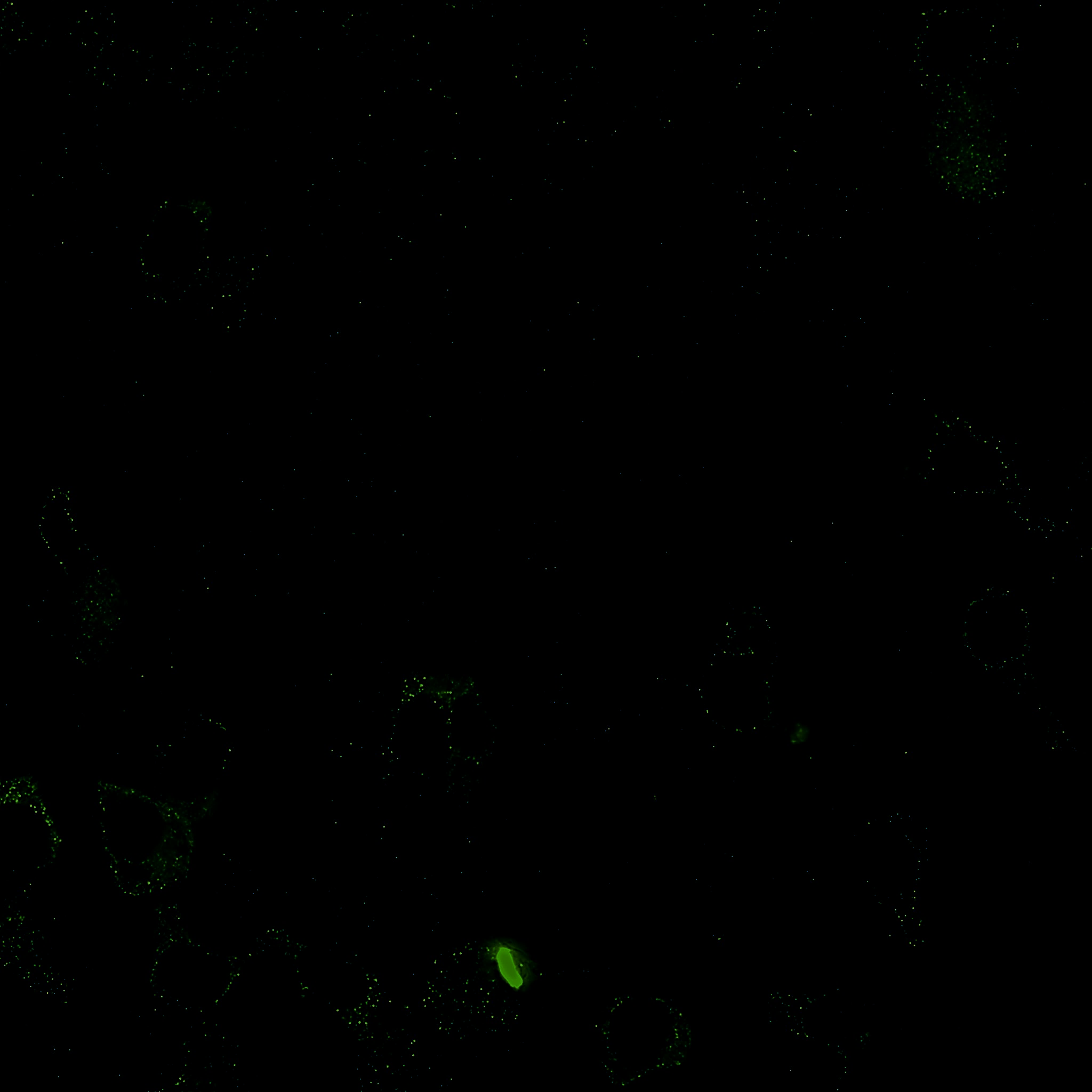

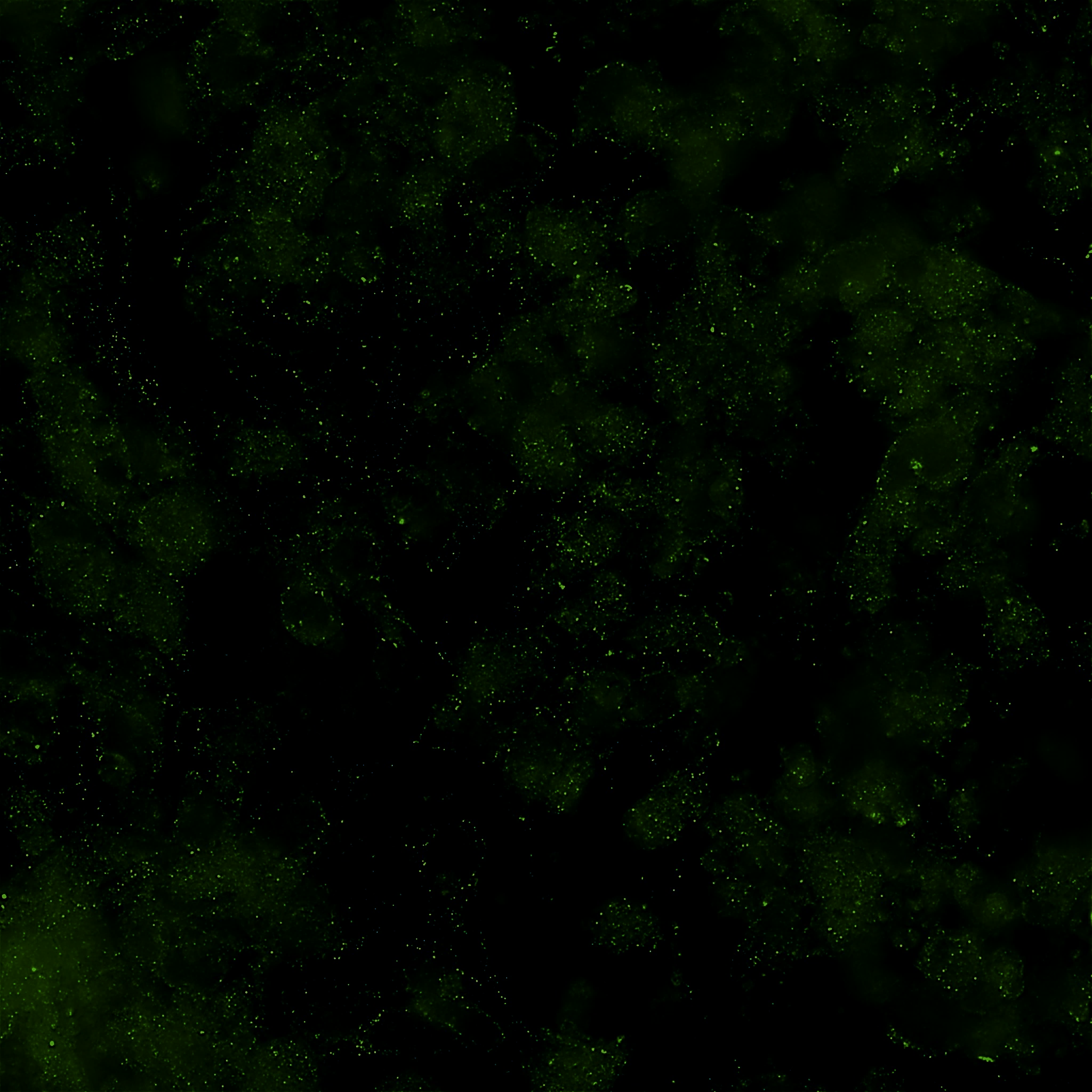

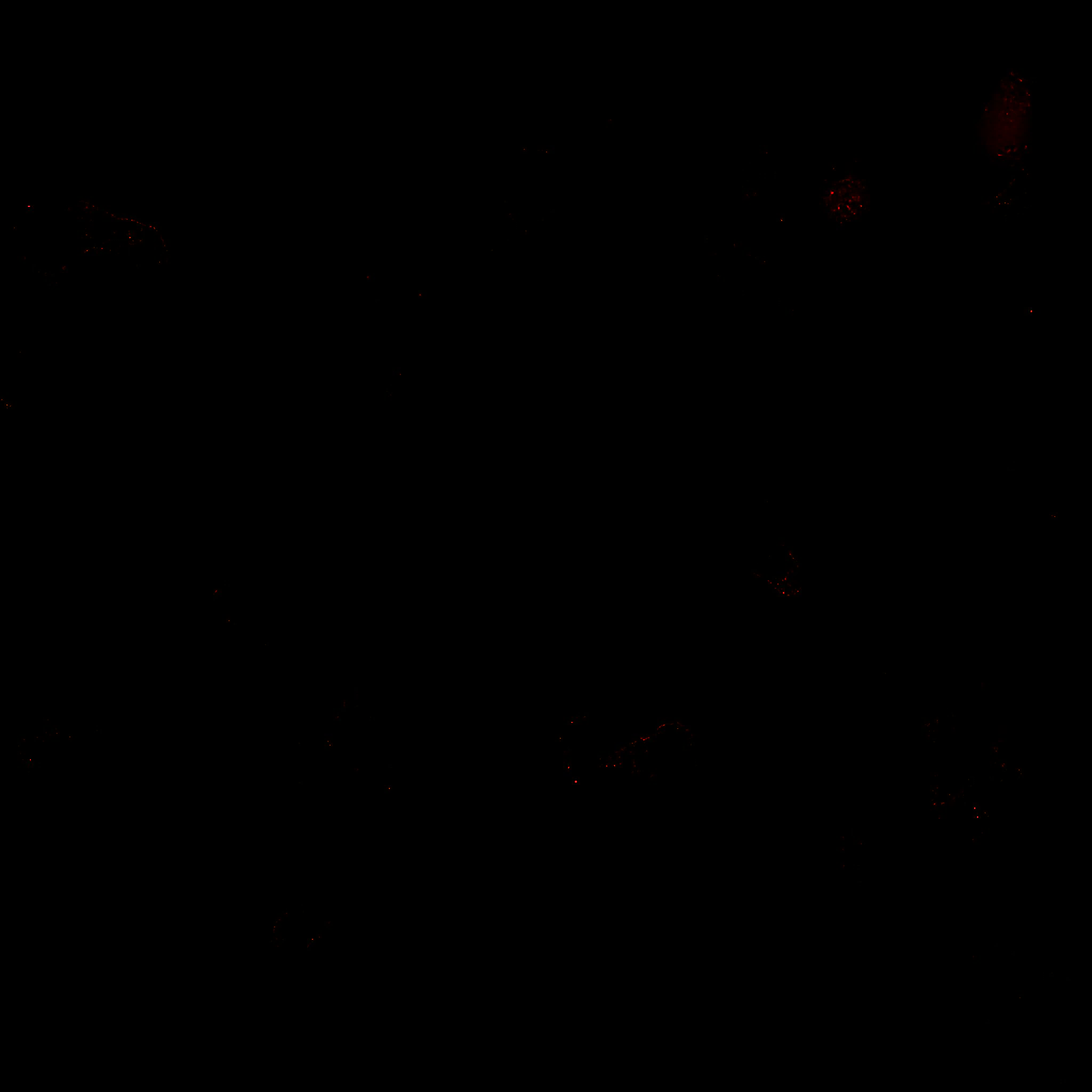

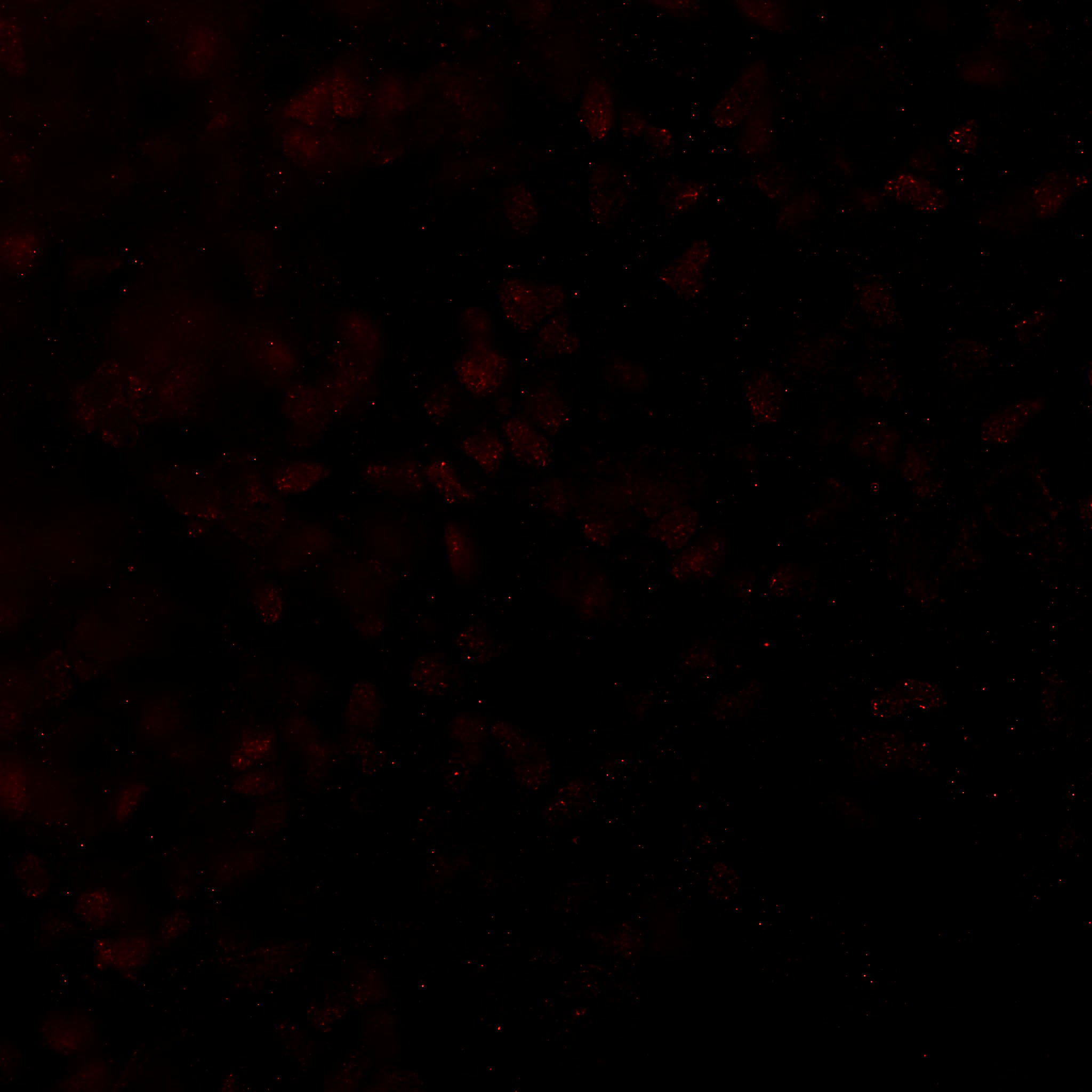

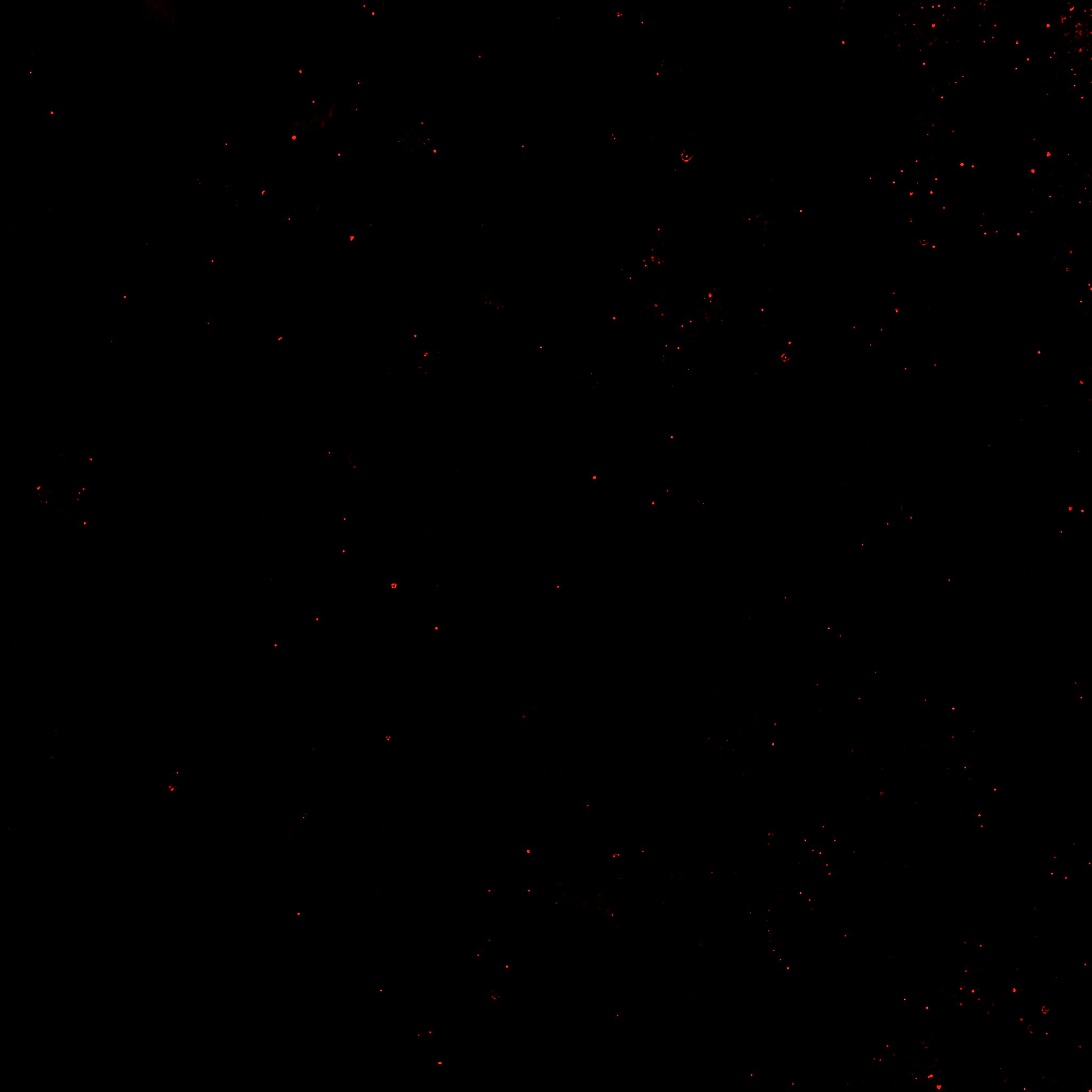

**PAX6**

**SOX17**

**NCAM**

**hESCs**

**Endoderm**

**Mesoderm**

**Ectoderm**

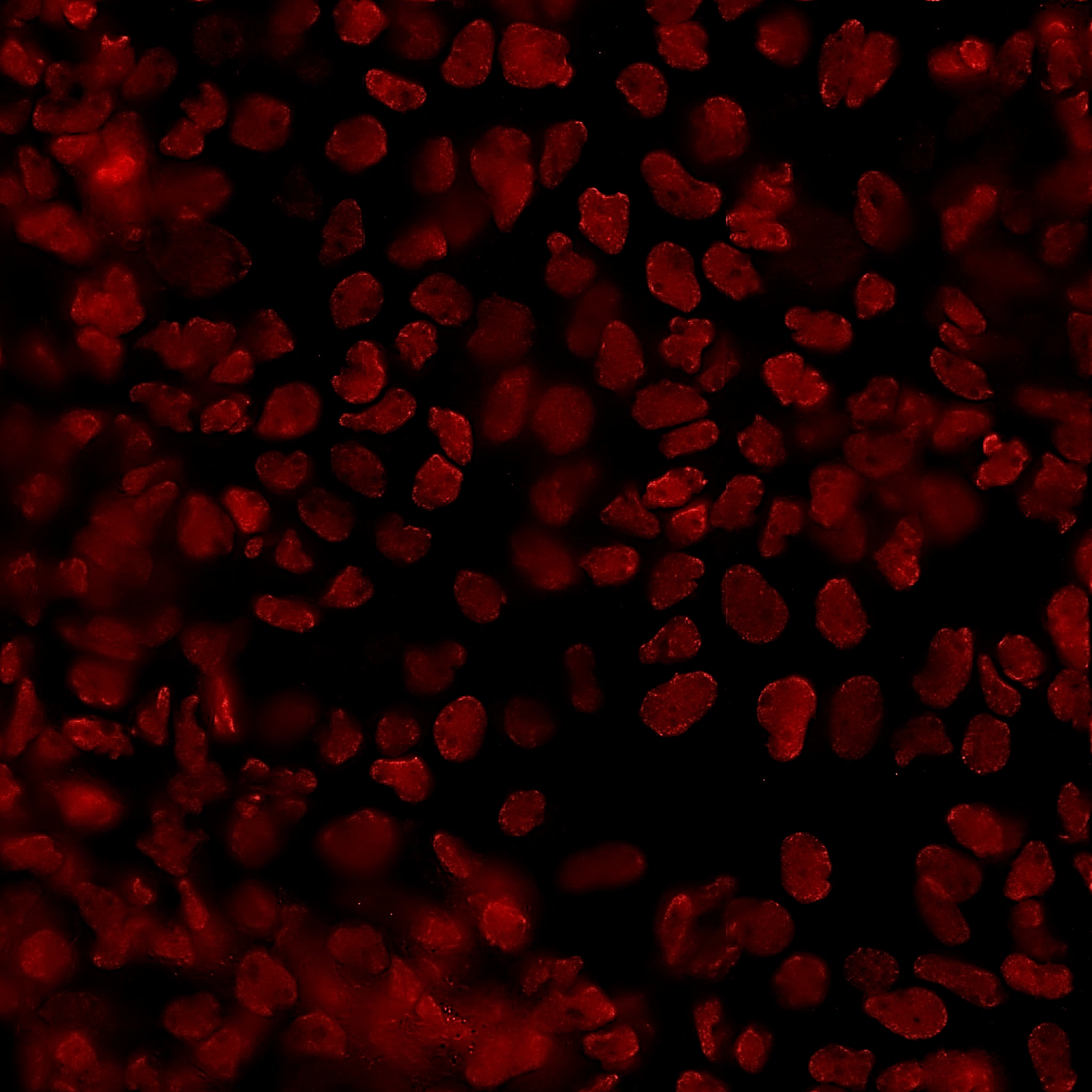

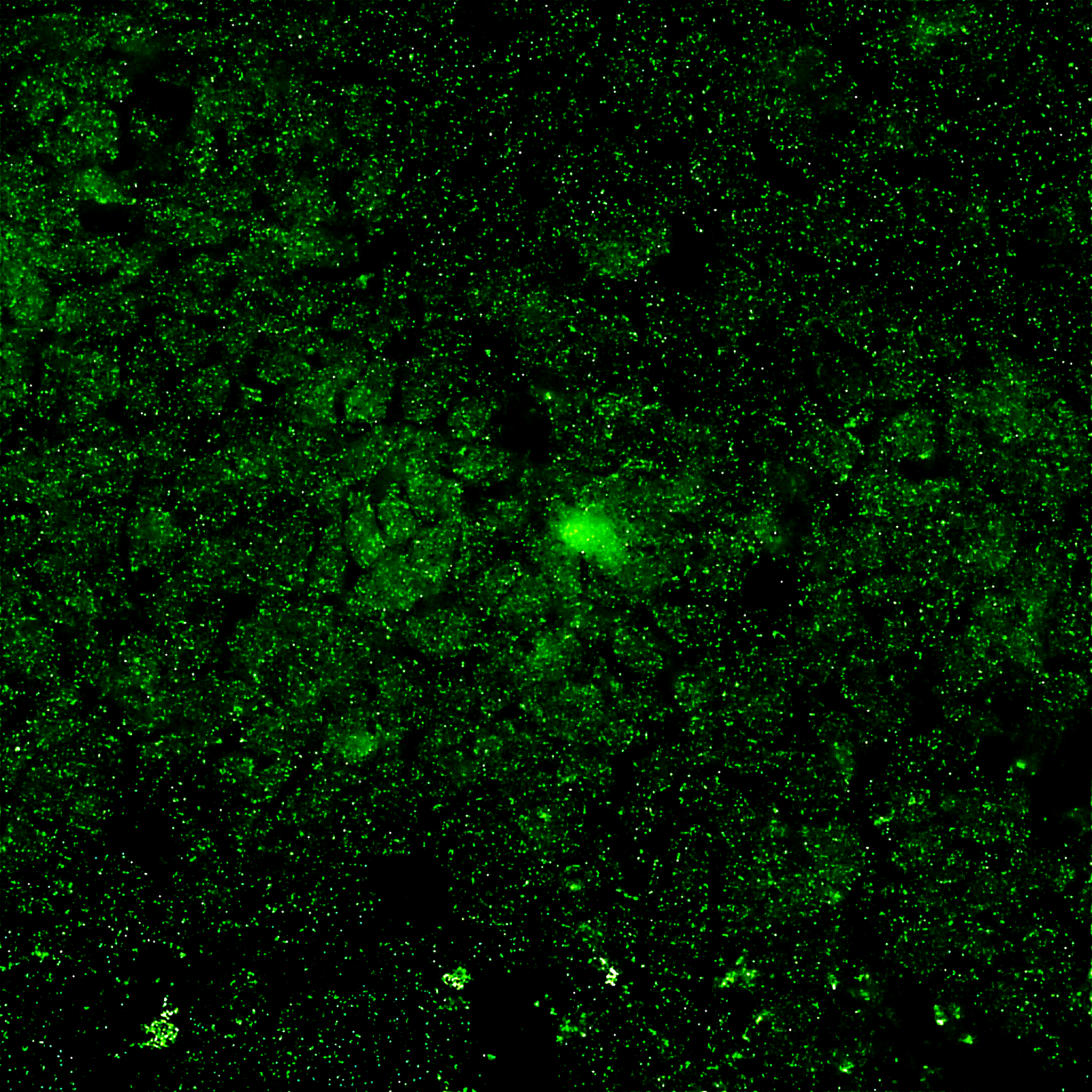

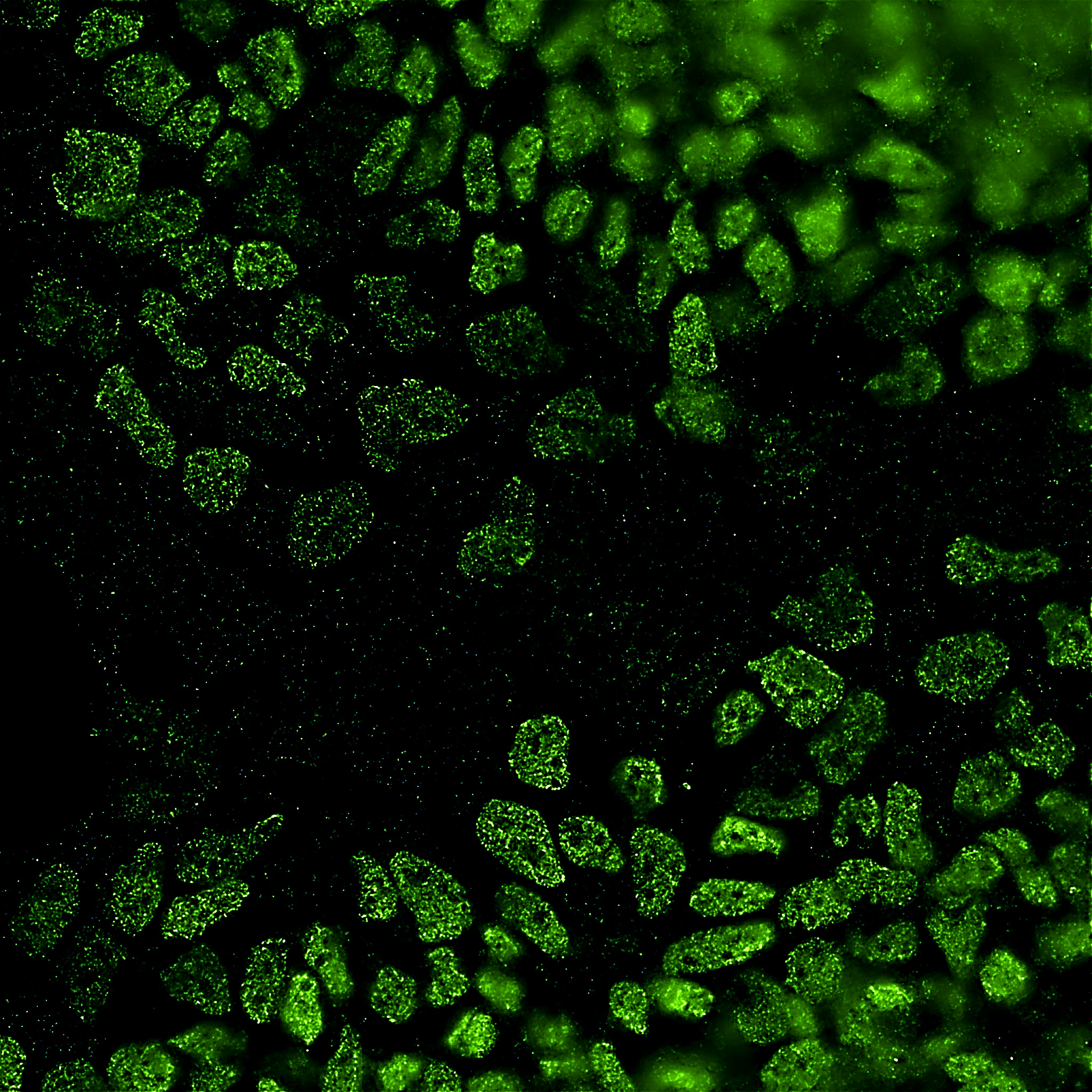

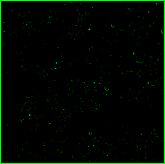

50 µm

Figure S1. The three germ layers were differentiated from H9 hESCs. Immunocytochemistry showing successful differentiation of H9 hESCs into endoderm, mesoderm, and ectoderm.

**Figure S2**

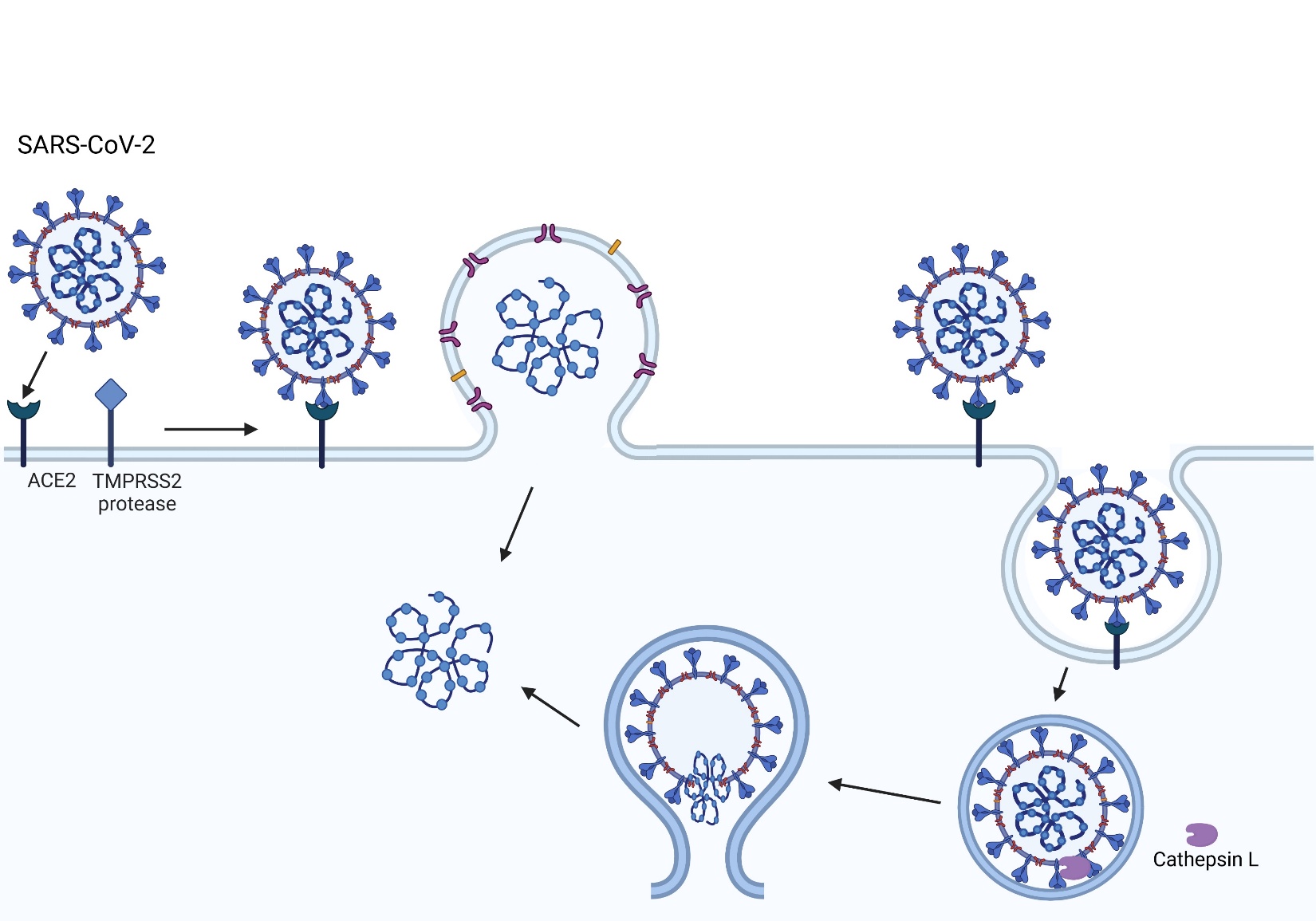

**A**

**B**

Figure S2. SARS-CoV-2 viral entry mechanisms. The SARS-CoV-2 virus binds to the ACE2 host cell receptor then either (A) TMPRSS2 processing of the viral spike protein leads to the fusion of the virus to the cell membrane or (B) the virus is taken up by endocytosis.

**Figure S3**

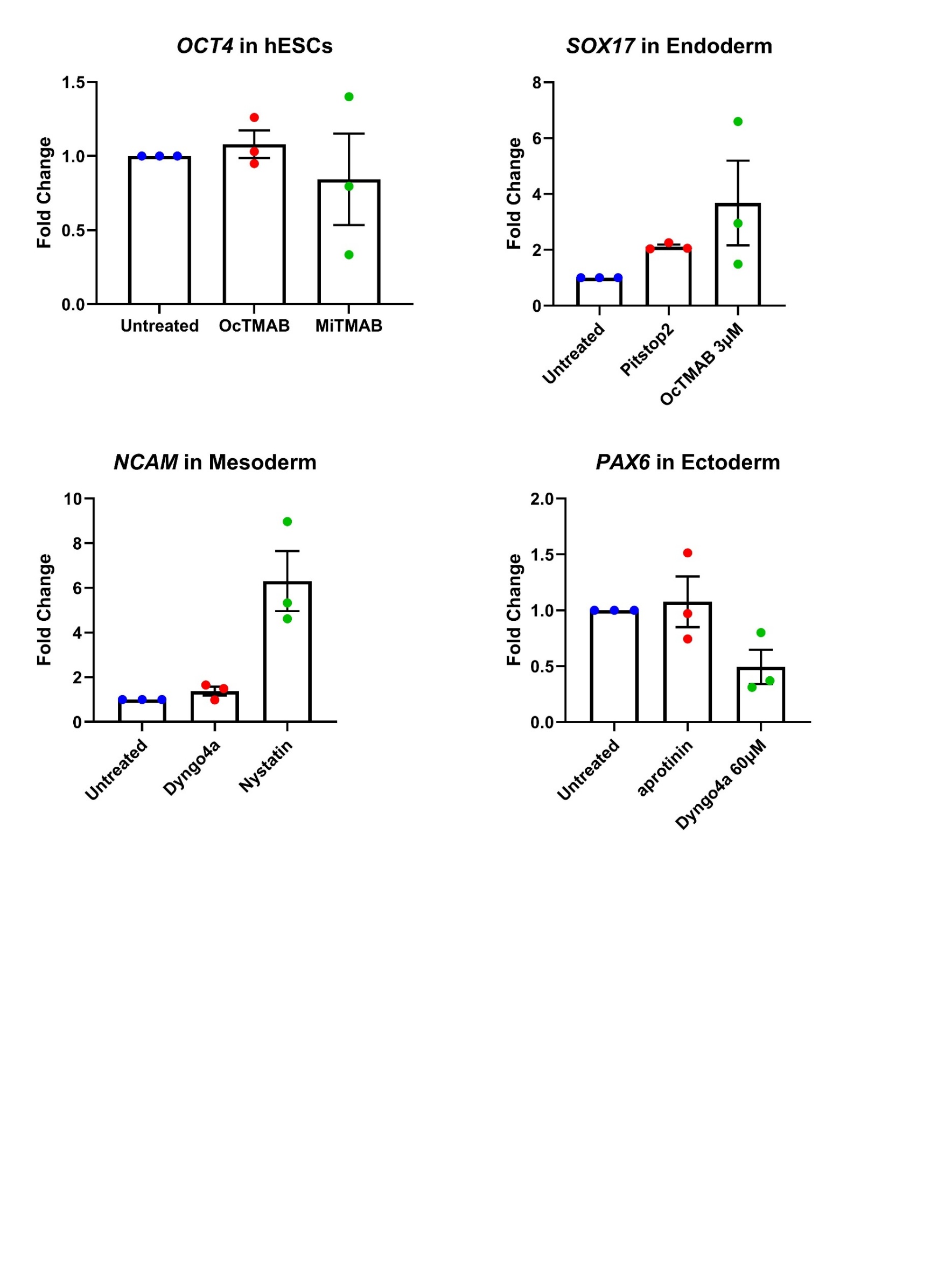

**A**

**B**

**C**

**D**

*****

*****

Figure S3. Most inhibitors did not significantly alter the expression of the hESC and germ layer markers. qPCR of (A) hESCs, (B) endoderm, (C) mesoderm, and (D) ectoderm was performed in the presence or absence of small molecule inhibitors to confirm the equivalent expression of markers for each cell type in control and treated groups. Kruskal-Wallis non-parametric analyses were performed on the qPCR data. Data are the means ± SEM of three independent experiments. * = p < 0.05.

### Table S1

| Name | Sequence |
| --- | --- |
| OCT4_F | CCTGAAGCAGAAGAGGATCACC |
| OCT4_R | AAAGCGGCAGATGGTCGTTTGG |
| PAX6_F | CTGAGGAATCAGAGAAGACAGGC |
| PAX6_R | ATGGAGCCAGATGTGAAGGAGG |
| SOX17_F | ACGCTTTCATGGTGTGGGCTAAG |
| SOX17_R | GTCAGCGCCTTCCACGACTTG |
| NCAM_F | CATCACCTGGAGGACTTCTACC |
| NCAM_R | CAGTGTACTGGATGCTCTTCAGG |
| β-actin_F | CACCATTGGCAATGAGCGGTTC |
| β-actin_R | AGGTCTTTGCGGATGTCCACGT |

**Table 1.** Name and sequences of qPCR primers.

**Figure S4**

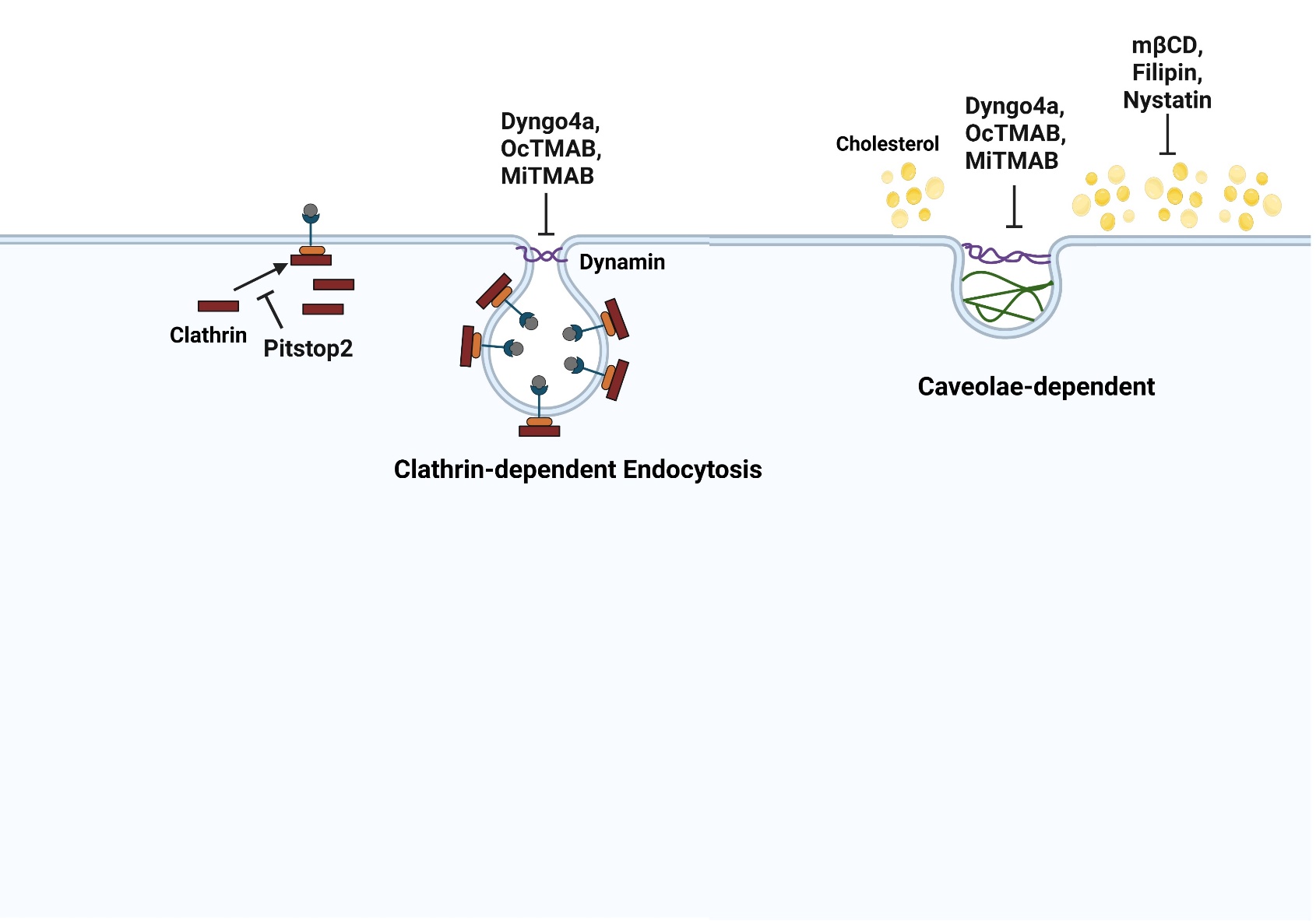

Figure S4. Diagram summarizing the effects of endocytosis inhibitors on SARS-CoV-2 entry. Dynamin-dependent endocytosis includes both clathrin- and caveolae- mediated endocytosis. The clathrin-mediated endocytosis pathway can be blocked by targeting various intermediates. Pitstop2 inhibits clathrin recruitment to the receptor. OcTMAB and MiTMAB blocks dynamin GTPase recruitment and dynamin activity is reduced by Dyngo4a. Caveolae-dependent endocytosis can be blocked by targeting cholesterol. mβCD, filipin, and genistein were moderately effective. mβCD and nystatin depletes cholesterol from the plasma membrane. Filipin loosens the packing of acyl chains by interacting with cholesterol.
